# Structural analyses of gibberellin-mediated DELLA protein degradation

**DOI:** 10.1101/2025.02.08.637281

**Authors:** Soyaab Islam, KunWoong Park, Eunju Kwon, Dong Young Kim

## Abstract

Gibberellin promotes plant growth by downregulating growth-repressor DELLA proteins. The gibberellin receptor GID1 binds to DELLA proteins in the presence of gibberellin, triggering their degradation through polyubiquitination by SCF^SLY1/GID2^ ubiquitin E3 ligase. Despite extensive studies, the molecular mechanisms by which DELLA proteins assemble with SCF^SLY1/GID2^ to regulate plant growth remain poorly understood. Here, we present two cryo-electron microscopy structures of the *Arabidopsis thaliana* DELLA protein RGA in complex with GID1A and GID1A-SLY1-ASK2, respectively. Structural analysis revealed that RGA interacts with GID1A and SLY1 through non-overlapping binding surfaces, stabilizing the proteins. This suggests that the SCF^SLY1^-RGA-GID1A complex assembles through stepwise stabilization induced by gibberellin. Furthermore, the structures indicate that RGA does not interact with IDD family transcription factors when bound to SLY1, suggesting that the binding of DELLA proteins to GID1/SLY1 and transcription factors is mutually exclusive. These findings provide insights into how DELLA proteins regulate transcription factor activity in response to gibberellin.

## INTRODUCTION

Gibberellin (GA) is a plant hormone that regulates various aspects of plant growth and development, including seed germination, stem elongation, flowering, and fertility. This hormone belongs to a group of structurally related diterpenoid acids, with a total of 136 natural compounds being identified as GAs in bacteria, fungi, and plants^1^. Among these, four GAs (GA_1_, GA_3_, GA_4_, and GA_7_) are considered bioactive forms^2^, which are perceived by the GA INSENSITIVE DWARF1 (GID1)^3–5^. GID1 is a soluble GA receptor composed of an N-terminal lid and a catalytically inactive hydrolase domain (IHD)^6,7^. The catalytically nonfunctional active site of the IHD forms a GA binding pocket, and the N-terminal lid closes the pocket through a disorder-to-order conformational change upon GA binding^6,7^. GA-bound GID1 interacts with DELLA proteins (named after the Asp-Glu-Leu-Leu-Ala sequence motif) to promote plant growth.

The DELLA proteins serve as coactivators or corepressors for various gene regulatory proteins that govern plant growth and development^8–11^. Several transcription factors featuring a basic helix-loop-helix (bHLH) DNA binding motif, such as PHYTOCHROME INTERACTING FACTORs (PIFs) and BRASSINAZOLE RESISTANT 1 (BZR1), bind to DELLA proteins and become inactivated^12–16^. Conversely, some INDETERMINATE DOMAIN (IDD) and basic leucine-zipper (bZIP) family proteins are activated upon interaction with DELLA proteins^17–21^. Physiologically, DELLA proteins act as repressors of plant growth, a function regulated by GA^8,9,22^. A depletion of GA, resulting from mutations in GA biosynthesis, leads to failed germination, extreme dwarfism, and unstable flowering development. The phenotypes associated with GA depletion can be rescued by an additional deletion mutation of the *DELLA* genes, indicating that the primary role of GA is to suppress the activity of DELLA proteins^8,9,22^. In this context, DELLA mutants with reduced GA sensitivity enable a “green revolution” wherein plants exhibit dwarfism while grain yields increase^23^.

DELLA proteins consist of an N-terminal regulatory domain (RD) and a C-terminal GRAS domain (named after the proteins GAI, RGA, and SCARECROW)^8,9^. The RD is responsible for the GA-dependent binding of DELLA proteins to GID1. The RD directly binds to the closed lid of GID1 upon GA binding through the DELLA, LExLE, and VHYNP motifs^6^. Consequently, the deletion of these motifs renders plant growth insensitive to GA^10,24^. The GRAS domain plays a crucial role in the transcriptional regulation of DELLA proteins; mutations in this domain result in the loss of DELLA protein activity, leading to a tall and slender growth phenotype^9,22,25^. While monocots such as rice and barley possess a single DELLA gene^9,25^, the model dicot *Arabidopsis thaliana* contains five *DELLA* genes: *REPRESSOR OF GA1-3* (*RGA)*, *GA-INSENSITIVE* (*GAI*), *RGA-LIKE1* (*RGL1*), *RGL2*, and *RGL3*^26^. Although these *DELLA* genes exhibit both distinct and redundant roles in regulating plant growth and development^27–29^, DELLA proteins are physiologically complementary to one another^14^, suggesting that the functional diversification of *DELLA* genes may result from their spatial and temporal expression patterns.

DELLA proteins are downregulated through polyubiquitination mediated by SLEEPY 1 (SLY1) in *Arabidopsis* and GID2 in rice. SLY1/GID2 acts as a substrate adaptor that recruits the GID1-GA-DELLA complex to the SCF (an acronym for SKP, CULLIN, F-BOX) ubiquitin E3 ligase for polyubiquitination and subsequent proteasomal degradation of DELLA proteins. Mutations in the *SLY1*/*GID2* gene block GA-induced proteolysis of DELLA proteins, resulting in a GA-insensitive phenotype^30–32^. This underscores the critical role of SCF^SLY1/GID2^-mediated degradation of DELLA proteins as a key regulatory pathway in GA-induced plant growth.

The physiological functions of DELLA proteins have been extensively studied. However, the molecular mechanisms by which GA regulates the interactions of DELLA proteins with SCF^SLY^^1/GID2^ and transcription factors remain poorly understood. This manuscript presents the cryogenic electron microscopy (cryo-EM) structures of the GID1A-GA_3_-RGA and GID1A-GA_3_-RGA-SLY1-ASK2 complexes, which were determined at resolutions of 2.66 Å and 2.80 Å, respectively. These structures provide insights into the mechanisms by which GA suppresses the activity of DELLA proteins and by which SCF^SLY1/GID2^ mediates their degradation.

## RESULTS

### Overall structure of the GID1A-RGA complex

To elucidate the interaction between GID1 and DELLA proteins, we purified the GID1A-RGA complex and determined its cryo-EM structure. The RGA protein, when expressed in *Escherichia coli*, was largely insoluble, with only a small soluble fraction that was unstable and prone to forming aggregates (**Fig. S1A–S1C**). GID1A was also mostly insoluble when expressed in *E. coli* in the presence of GA_3_, but small soluble fractions were eluted in a homogeneous form through size-exclusion chromatography (SEC) (**Fig. S1D–S1F**). The solubility and homogeneity of RGA and GID1A significantly improved when they were coexpressed in the presence of GA_3_. The GID1A-RGA complex, expressed in the presence of GA_3_, was purified via immobilized metal affinity chromatography (IMAC) and SEC **(Fig S1G–S1I)**. The SEC-multiangle light scattering (MALS) experiment demonstrated that the purified GID1A-RGA forms a homogeneous heterodimer (**Fig. 1A**). The cryo-EM map of the GID1A-RGA complex was reconstructed at an overall resolution of 2.66 Å, using 251,864 particles selected from the cryo-EM images of the purified GID1A-RGA (**Figs. 1B–D, S2, and S3**). The atomic model was built by tracing the cryo-EM map and refining it through iterative model correction and structural refinement. The correlation coefficient between the final structure and map (CC_mask_) was 0.84, and no outliers were found in the Ramachandran plot. The final structure comprised one GID1A monomer, one RGA monomer, one GA_3_ molecule, and 248 water molecules (**Fig. S3D and Table S1**).

**FIGURE 1.**
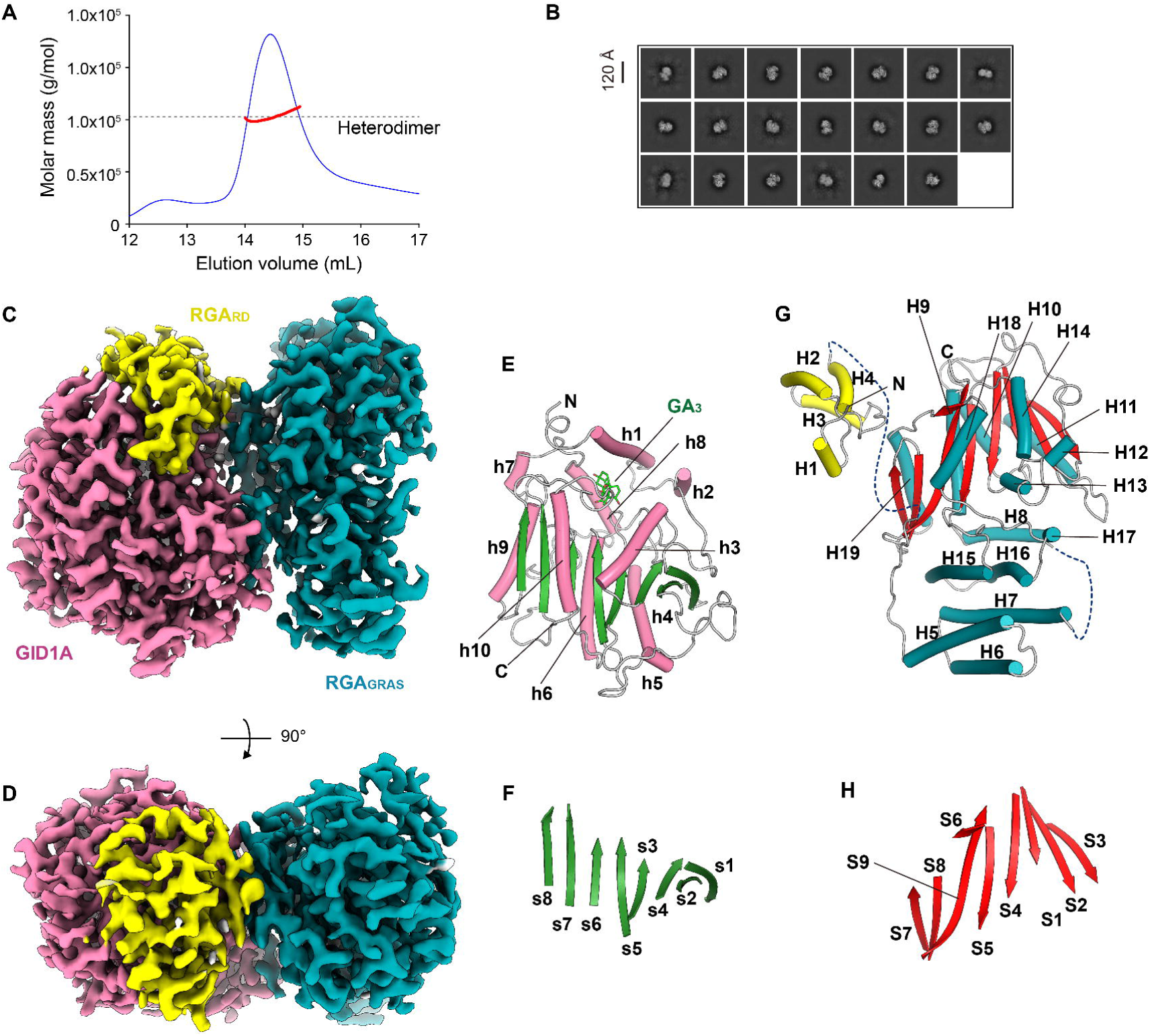
Cryo-EM structure of the RGA-GID1A complex. (A) SEC-MALS analysis of purified RGA-GID1A. The absorbance at 280 nm is shown in blue, and the experimental molar masses of the eluate are represented in red. The dotted line indicates the calculated molecular weight of the RGA-GID1A heterodimer (102.6 kDa). (B) Representative two-dimensional classes of RGA-GID1A particles. (C, D) Cryo-EM map of RGA-GID1A shown in two different orientations. GID1A, RGA_RD_, and RGA_GRAS_ are colored pink, yellow, and cyan, respectively. (E, F) GA_3_-bound GID1A within the cryo-EM structure of RGA-GID1A. The central β-sheet is highlighted in (F). The secondary structures are denoted as h1–h10 for the helices and s1–s8 for the strands. GA_3_ is depicted as a green stick model. (G, H) RGA in the cryo-EM structure of RGA-GID1A. The core β-sheet is shown in (H). The secondary structure is denoted as H1–H19 for helices and S1–S9 for strands. The helices in (E) and (G) are color-coded as in the cryo-EM maps shown in (C) and (D).

In the cryo-EM structure of the GID1A-RGA complex, GID1A consists of an N-terminal lid (GID1A_Lid_; residues 10–50) and an IHD (GID1A_IHD_; residues 61–343) (**Fig. 1E**). GID1A_IHD_ exhibited an α/β fold with a parallel β-sheet (topological order: s1, s2, s4, s3, s5, s6, s7, and s8) and peripheral helices (h3–h10) (**Figs. 1E, 1F, and S4A**). GID1A_Lid_ contains two helices (h1 and h2) that cover the GA_3_ molecule bound to GID1A_IHD_, completely enclosing GA_3_ within the space between GID1A_IHD_ and GID1A_Lid_. The overall fold of GID1A is nearly identical to that observed in the crystal structure of the GID1A-GAI_RD_ complex (PDB ID: 2ZSH)^6^ (**Fig. S4B**), with an RMSD value of 0.6 Å for 334 Cα atoms.

RGA comprises an N-terminal RD (RGA_RD_; residues 42–108) and a C-terminal GRAS domain (RGA_GRAS_; residues 205–584) (**Fig. 1G**). The segment (residues 109–204) linking RGA_RD_ and RGA_GRAS_ was not visible in the cryo-EM map. The flexible linker among DELLA proteins was highly variable in both length and sequence (**Fig. S5**), suggesting that this linker may play a role in the specific functions of each DELLA protein. RGA_RD_ consists of four helices (H1–H4) and forms a concave groove that facilitates GID1A binding (**Fig. 1F**). The RGA_RD_ region in the cryo-EM structure did not exhibit significant conformational differences when compared to GAI_RD_ in the crystal structure of the GAI_RD_-GID1A complex (PDB ID: 2ZSH), with an RMSD value between RGA_RD_ and GAI_RD_ of 1.3 Å for 60 Cα atoms (**Fig. S6A**). RGA_GRAS_ displayed an α/β fold with a twisted core β-sheet and peripheral helices (H8–H14, H18, and H19) (**Fig. 1G**). The core β-sheet consisted of eight β-strands arranged topologically in the order S3, S2, S1, S4, S5, S9, S8, and S7 (**Fig. 1H**). Strand S9 is longer than the other strands and forms additional hydrogen bonds with strand S6, aligning in the same direction as strand S8 and extending beyond the core β-sheet. In addition to the α/β fold, RGA_GRAS_ features a helical bundle composed of helices H5–H8, H15, and H16. Overall, RGA_GRAS_ exhibited a rod-shaped structure that included both the α/β fold and the helical bundle. A structural homology search via the DALI server^33^ indicated that RGA_GRAS_ shares a fold with GRAS family proteins. Notably, *Arabidopsis* SCARECROW-LIKE 3 (SCL3; PDB ID: 6KPD) was superimposed onto RGA_GRAS_, yielding the lowest RMSD value of 1.7 Å for 353 Cα atoms (**Fig. S6B**). Some GRAS domains form homodimers or heterodimers through their helical bundles, corresponding to the LHRI region^34,35^. It has been suggested that RGA also forms a homodimer through its LHRI region (helices H5–H7)^15^. However, RGA_GRAS_ in the cryo-EM structure of GID1A-RGA did not form a homodimer despite the solvent-exposed surface of the LHRI region (**Fig. S6C and S6D**). Consistent with this observation, larger particles containing RGA homodimers were not detected in the two-dimensional (2D) classification of the cryo-EM particles (**Fig. S2**). The surfaces of GID1A and RGA_RD_ at their binding interface are partially positively and negatively charged, respectively. Conversely, those of GID1A and RGA_GRAS_ at their binding interface are relatively hydrophobic (**Fig. S6E and S6F**).

### The binding of GA_3_ in the GID1A pocket

GID1A_IHD_ possesses a GA-binding pocket formed by the helices h3 and h8, the preceding loops (l2 and l3), and the loop l4 located between strand s8 and helix h10 (**Fig. 2A and S4B**). GA_3_ directly forms hydrogen bonds with residues S116, Y127, and F238 of GID1A_IHD_, and hydrophobic interaction with Y247 through its methyl group within the pocket (**Fig. 2A**). The GA-binding pocket was slightly larger than GA_3_, allowing water molecules to occupy the space and mediate interactions between GA_3_ and the pocket (**Fig. 2A**). In the crystal structure of the GAI_RD_-GID1A complex^6^, four water molecules bridged the interactions between GID1A and GA_3_. In the cryo-EM structure of the RGA-GID1A complex, three water molecules directly mediate the interactions between GID1A and GA_3,_ similar to the interactions observed in the crystal structure of GAI_RD_-GID1A (**Fig. 2A**). The GA-binding pocket was directly covered by helix h1 of GID1A_Lid_ (**Fig. 2B**). GA_3_ bound F27 in helix h1 of GID1A_Lid_ through hydrophobic interaction. As with the GA_3_-binding pocket, water molecules also mediate additional interactions between GA_3_ and GID1A_Lid_ (**Fig. 2B and 2C**).

**FIGURE 2.**
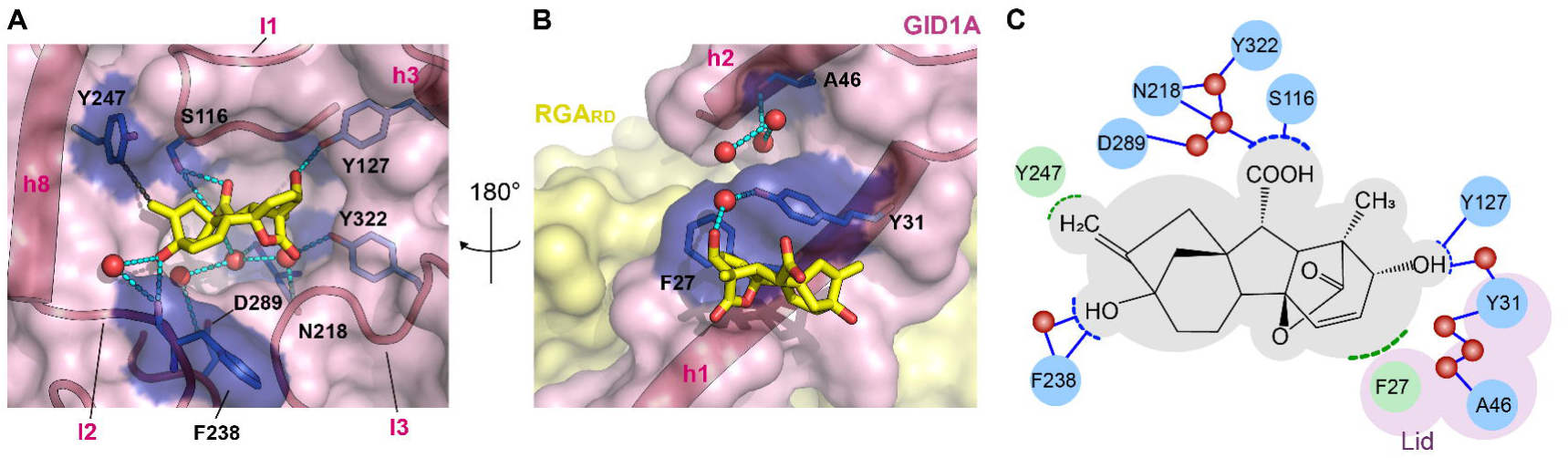
The GA_3_-binding pocket of GID1A. (A) GA_3_ in the binding pocket. (B) GA_3_ covered by GID1A_Lid_. (A) and (B) are views from opposite directions centered on GA_3_. GA_3_ is shown as a yellow stick model, and GID1A is represented as a pink surface model. Residues interacting with GA_3_ are depicted as blue stick models and surface models. Water molecules are shown as red spheres, and the hydrogen bonding network around GA_3_ is indicated by dotted lines. The yellow surface indicates the RGA. (C) Interactions between GID1A and GA_3_. The GID1A residues involved in the hydrogen bonding network and hydrophobic interactions are highlighted in blue and green, respectively.

### Interactions between GID1A and RGA

The RGA interacted with GA_3_-bound GID1A over an extensive binding interface that encompassed both RGA_RD_ and RGA_GRAS_ (**Fig. 3A**). The RGA surface area was buried by 11.4% within the GID1A-binding interface (2,498 Å^2^ in 21,833 Å^2^), resulting in a reduction in ΔG by −33.2 kcal/mol. Among the four helices of RGA_RD_, helix H1, which contains the DELLA motif, contacts the surface around helix h3 of GID1A mainly through hydrophobic interactions. Additionally, a water molecule bridged interactions between RGA_RD_-L46 and GID1A-R133/L323 through hydrogen bonding (**Fig. 3B**). Helix H2 of RGA_RD_ contacted the N-terminal loop and helix h1 of GID1A, which encapsulates GA_3_. RGA_RD_-E67 formed ionic bonds with GID1A-R13, and RGA_RD_-E70 formed hydrogen bonds with GID1A-K28, which is the next residue of F27 that directly contacts GA_3_ (**Fig. 3C**). Residues participating in hydrophobic interactions were clustered near the ionic and hydrogen bonds. Helix H3 of RGA_RD_ contacted the C-terminus of helix h1. The loop L1 between helices H3 and H4 formed a slight contact with helix h2 (**Fig. 3D**). Helix H4, which contains the VHYNP motif, contacts helix h1 of GID1A through hydrophobic interactions. These observations indicate that RGA_RD_ is responsible for GID1A_Lid_ binding. The surface of RGA_RD_ is buried by 26.9% within the binding interface (1,310 Å^2^ in 4,869 Å^2^), contributing to a ΔG reduction of −22.8 kcal/mol.

**FIGURE 3.**
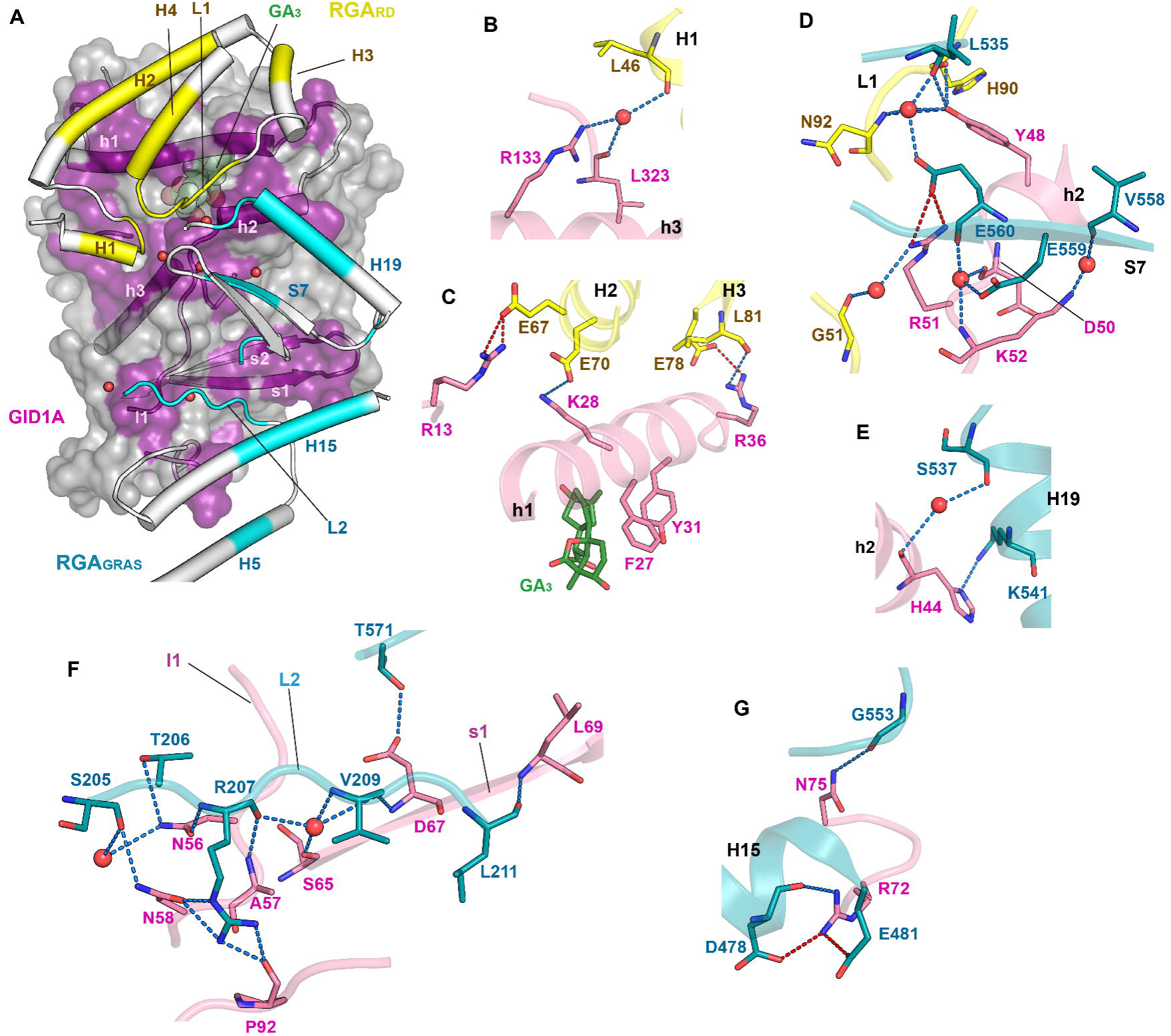
Interactions between GID1A and RGA. (A) The binding interface between GID1A and RGA. RGA is shown as a low-transparency surface model. RGA and GID1A at the binding interface are depicted as cartoon models. Residues of GID1A, RGA_RD_, and RGA_GRAS_, involved in interactions at the binding interface, are highlighted in purple, yellow, and cyan, respectively. Water molecules are shown as red spheres. (B–G) Detailed interactions between GID1A and RGA. The GID1A, RGA_RD_, and RGA_GRAS_ models are presented in pink, yellow, and cyan, respectively. The blue and red dotted lines indicate hydrogen bonds and ionic interactions, respectively. (B, C) Interactions between GID1A and RGA_RD_. (D) Interactions of GID1A with RGA_RD_ and RGA_GRAS_ at the junctional area. (E–G) Interactions between GID1A and RGA_GRAS_.

RGA_GRAS_ interacts with GID1A through multiple binding motifs, forming a large binding interface (**Fig. 3A**). First, helix H19 and strand S7 of RGA_GRAS_ contacted residues surrounding helix h2 of GID1A (**Fig. 3D and 3E**). RGA_GRAS_-E560 formed an ionic bond with GID1A-R51, and L535 and K541 of RGA_GRAS_ established hydrogen bonds with GID1A-Y48 and GID1A-H44, respectively. Notably, GID1A-Y48 interacted with both RGA_GRAS_ and RGA_RD_ (**Fig. 3D**). Second, loop L2 (residues 205–211) of RGA_GRAS_, which serves as a linker connecting RGA_RD_ and RGA_GRAS_, contacts the surface loops and strand s1 of GID1A. The residues S205, T206, R207, V209, and L211 in loop L2 of RGA_GRAS_ formed hydrogen bonds with the GID1A residues N58, N56, N56/A57/N58/P92, D67, and L69, respectively (**Fig. 3F**). Third, helix H15 of RGA_GRAS_ contacted the loop between strands s1 and s2. RGA_GRAS_-D478 and E481 formed ionic bonds with GID1A-R72 (**Fig. 3G**). Additionally, several hydrophobic interactions further contributed to the interaction between RGA_GRAS_ and GID1A. These hydrophobic interactions were predominantly found around hydrogen and ionic bonds, and helix H5 also bound to GID1A-L95 through hydrophobic interactions. The RGA_GRAS_ surface was buried by 7.2% within the binding interface (1,258 Å^2^ in 17,497 Å^2^), with a ΔG reduction of −10.7 kcal/mol. Furthermore, seven water molecules bridged the interactions between RGA_GRAS_ and GID1A (**Fig. 3B, 3D–F**). Despite the extensive binding area of RGA_GRAS_, GID1A, when coexpressed with RGA_GRAS_ in the presence of GA_3_, precipitated during purification under high-salt buffer conditions (**Fig. S7A–I**). This observation suggests that RGA_RD_ is a prerequisite for the interactions and stability between RGA_GRAS_ and GID1A.

### Overall structure of the GID1A-RGA-SLY1-ASK2 *complex*

DELLA proteins are polyubiquitinated by SCF^SLY1/GID2^ ubiquitin E3 ligase in the presence of GA, leading to their proteasomal degradation^30–32^. To elucidate the mechanisms by which the DELLA protein binds to SCF^SLY1/GID2^, we purified the *Arabidopsis thaliana* GID1A-RGA-SLY1-ASK2 complex and determined its cryo-EM structure. The GID1A-RGA-SLY1-ASK2 complex was purified via a copurification strategy. When SLY1 and ASK2 were coexpressed, only ASK2 was successfully purified, whereas SLY1 precipitated during purification due to its low stability (**Fig. S7K–L**). However, similar to the copurification of GID1A and RGA (**Fig. S1**), the stability of SLY1 was significantly enhanced when SLY1-ASK2 was co-purified with GID1A-RGA in the presence of GA_3_ (**Fig. S7M–O**). To purify the GID1A-RGA-SLY1-ASK2 complex, GID1A-RGA was expressed in *E. coli* in the presence of GA_3_, and the SLY1-ASK2 complex was expressed in *E. coli* without GA_3_. The cell lysates containing the recombinant GID1A-RGA and SLY1-ASK2 were mixed in balanced quantities. Purification of the GID1A-RGA-SLY1-ASK2 complex was performed via IMAC and SEC under high-salt buffer conditions with 0.5 M NaCl (**Fig. S7M–O**). The SEC-MALS experiment demonstrated that the purified GID1A-RGA-SLY1-ASK2 complex forms a homogeneous heterotetramer (**Fig. 4A**). These results indicate that the protein assembly of the GID1A-RGA-SLY1-ASK2 complex stabilizes the individual proteins involved. To determine the cryo-EM structure of the GID1A-RGA-SLY1-ASK2 complex, its cryo-EM map was reconstructed at 2.80 Å resolution, with 65,737 particles selected from cryo-EM images of the purified complex. As the cryo-EM map did not clearly show the density of SLY1-ASK2, an additional map focusing on SLY1-ASK2 was constructed at an overall resolution of 3.08 Å. The complete map covering all regions of the GID1A-RGA-SLY1-ASK2 complex was obtained by combining these two maps (**Figs. 4B–D, S8, and S9**). The final model structure of the GID1A-RGA-SLY1-ASK2 complex exhibited a CC_mask_ of 0.78, with no outliers in the Ramachandran plot (**Table S1**).

**FIGURE 4.**
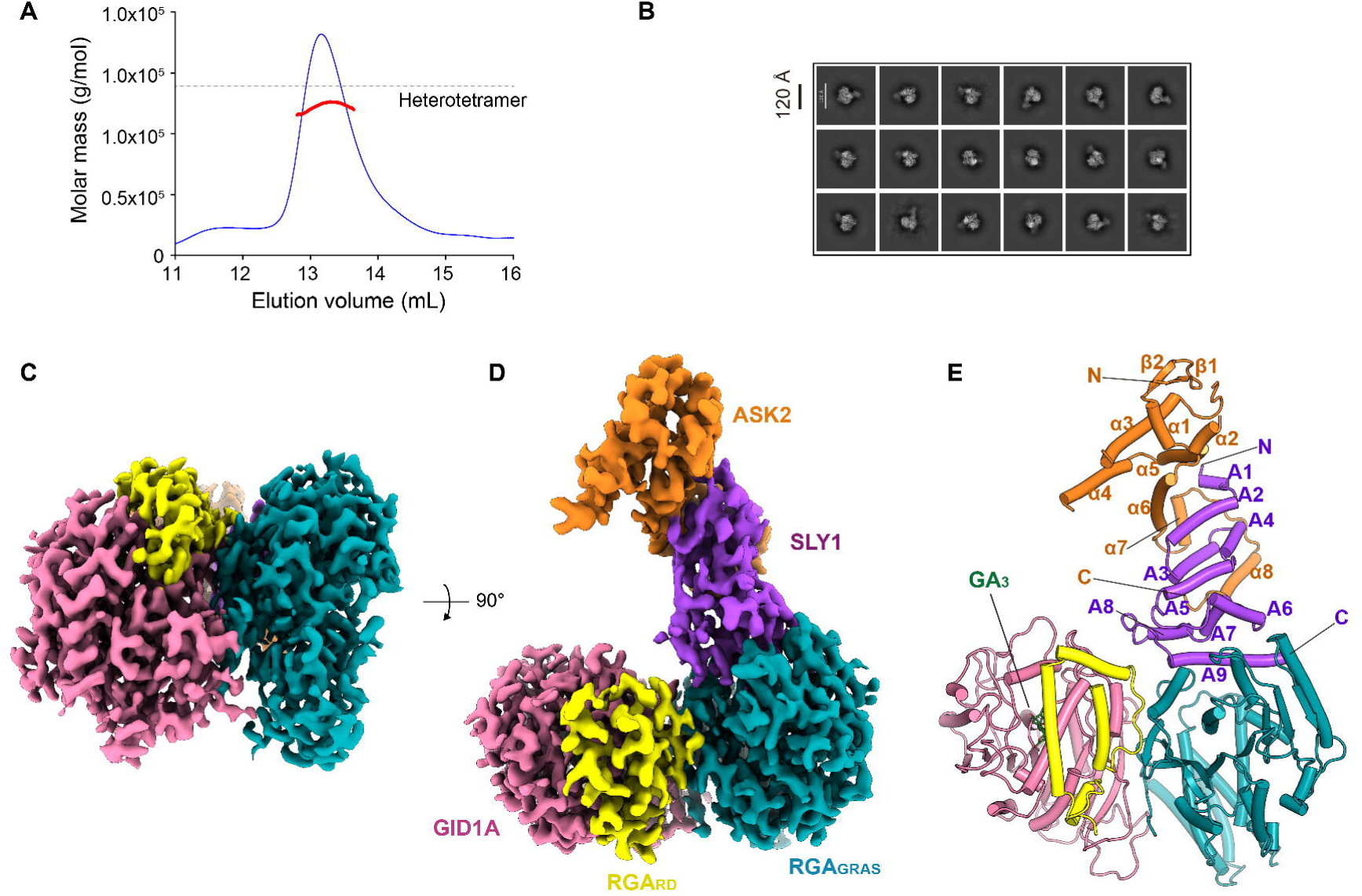
Cryo-EM structure of the GID1A-RGA-SLY1-ASK2 complex. (A) SEC-MALS analysis of purified RGA-GID1A-SLY1-ASK2. The absorbance at 280 nm is shown in blue, and the experimental molar masses of the eluate are depicted in red. The dotted line indicates the calculated molecular weight of the RGA-GID1A-SLY1-ASK2 heterotetramer (138.6 kDa). (B) Representative two-dimensional classes of RGA-GID1A-SLY1-ASK2 particles. (C, D) Cryo-EM map showing two different orientations. GID1A, RGA_RD_, RGA_GRAS_, SLY1, and ASK2 are colored pink, yellow, cyan, purple, and orange, respectively. (E) Cartoon model in the same orientation as (D). The color scheme matches the cryo-EM map in (C) and (D). The secondary structures are labeled A1–A9 for the helices of SLY1, α1–α8 for the helices of ASK2, and β1–β2 for the strands of ASK2.

The GID1A-RGA in the cryo-EM structure of GID1A-RGA-SLY1-ASK2 was nearly identical to that in the cryo-EM structure of GID1A-RGA. Both GID1A-RGA structures were superimposed, with an RMSD value of 0.6 Å for 773 Cα atoms (**Fig. S10A**). No significant conformational differences were detected between the two GID1A-RGA complexes, except for a local conformational change in the loop between strand S2 and helix H11 of RGA (residues 362–367). Although this loop is situated near the SLY1-binding interface of RGA, no direct interaction between the loop and SLY1 was observed (**Fig. 4D**).

SLY1 formed a helical structure comprising nine helices (Α1–A9) (**Figs. 4E and S10B**). The three N-terminal helices A2–A4 form an F-box motif responsible for SKP1 binding, and the C-terminal long helix A9 is oriented toward the RGA surface. Structural homology searches revealed that SLY1 shares an overall fold with human F-box-only protein 31 (FBXO31; PDB ID: 5VZT), except for helix A9. Superimposing SLY1 with FBXO31 yielded an RMSD value of 2.5 Å for 90 Cα atoms (**Fig. S10C**). In the cryo-EM structure, SLY1 binds to ASK2 and RGA through its N-terminal F-box motif and C-terminal helix A9, respectively, thereby linking RGA to the SCF complex as a substrate adaptor for the ubiquitin E3 ligase. ASK2 formed a helical structure consisting of seven helices (α1–α7) and contains an additional two-stranded short β-sheet at its N-terminus (**Fig. 4E**). ASK2 exhibited a fold similar to that of the F-box binding protein SKP1 of the SCF complex and it was superimposed with *Arabidopsis thaliana* ASK1, with the lowest RMSD value (1.2 Å for 132 Cα atoms).

### SLY1 as a substrate adaptor for RGA ubiquitination

In the cryo-EM structure of the GID1A-GA_3_-RGA-SLY1-ASK2 complex, RGA_GRAS_ forms a surface groove near RGA_RD_ that binds SLY1. Although RGA_RD_ and GID1A are located close to SLY1, no direct interaction was observed between SLY1 and either RGA_RD_ or GID1A (**Fig. 4B**). Helix A9 of SLY1 directly binds to helices H9–H11 of RGA_GRAS_. Specifically, residues K126, S130, Y137, and S141 of helix A9 in SLY1 formed hydrogen bonds with residues Q311, A338/Q341, P337, Q332, and K375 of RGA_GRAS_, respectively (**Fig. 5A**). Additionally, the RGA-binding interface of SLY1 exhibited high hydrophobicity (**Fig. 5B and 5C**). Twenty-two hydrophobic interactions were identified around the hydrogen bonds between RGA_GRAS_ and SLY1, suggesting that the hydrophobic surface of SLY1 is stabilized through binding to RGA. This is consistent with the observation that SLY1 precipitated during the purification of the SLY1-ASK2 complex but was stabilized and coeluted during copurification with ASK2 and GID1A-RGA (**Fig. S7**). Moreover, SLY1 was stabilized and coeluted with ASK2 and RGA_GRAS_ in SEC during purification when SLY1-ASK2 was copurified with RGA_GRAS_ (**Fig. S7P–R**). These findings suggest that SLY1 stability is enhanced through its interactions with RGA_GRAS_ and ASK2.

**FIGURE 5.**
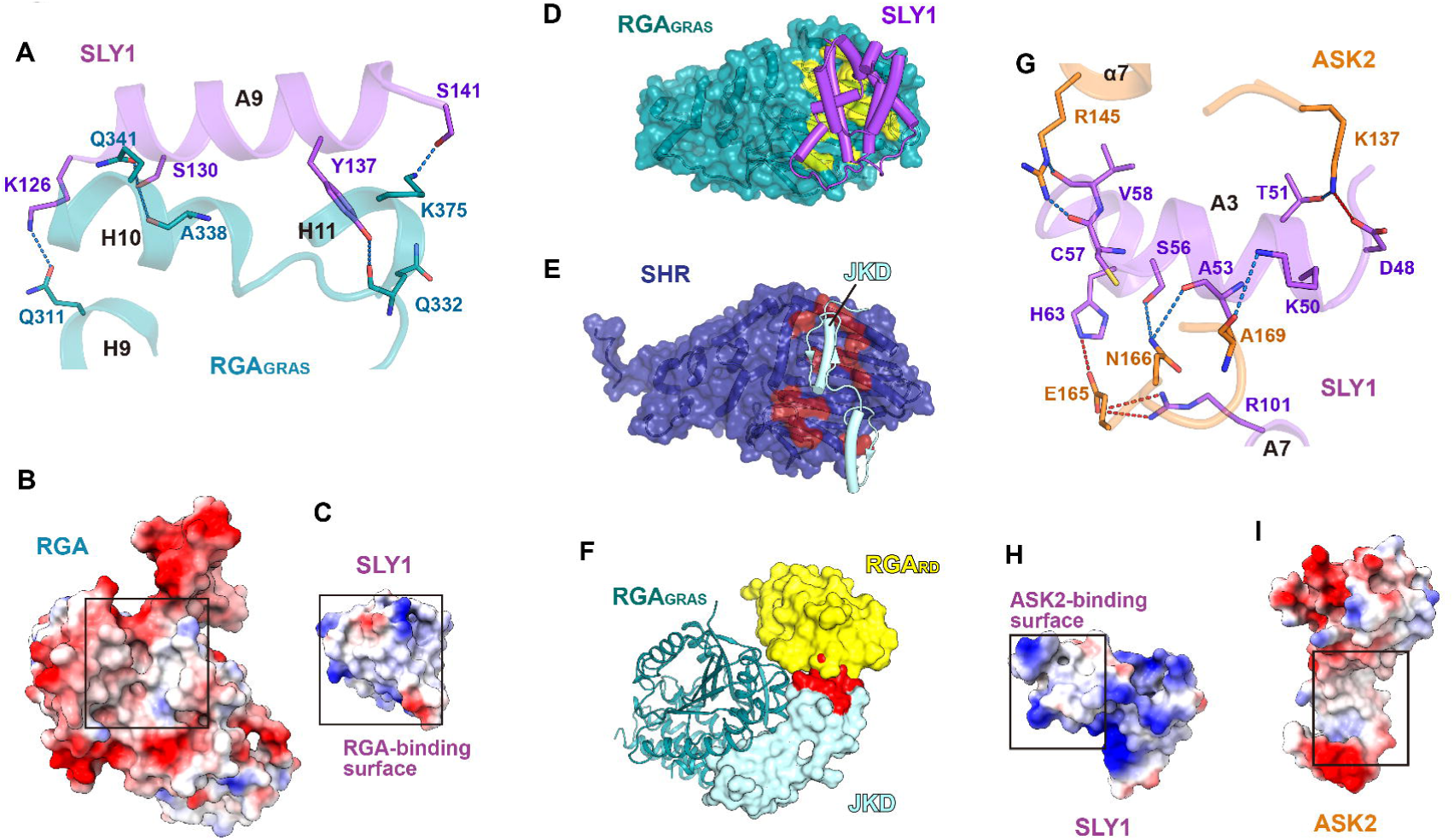
Interactions of SLY1 with RGA_GRAS_ and ASK2. (A) Cartoon model depicting the binding interface between SLY1 and RGA_GRAS_. The residues involved in the interactions are shown as purple and cyan stick models for SLY1 and RGA_GRAS_, respectively. (B, C) Surface charge distribution at the binding interface of RGA_GRAS_ (B) and SLY1 (C). The black-lined boxes highlight the binding interface. The electrostatic potential from red (−10 kcal/mol·*e*) to blue (+10 kcal/mol·*e*) is mapped onto the solvent-accessible surfaces. (D, E) Structural comparison between RGA_GRAS_-SLY1 (D) and SHR-JKD (E). RGA_GRAS_ and SHR are shown in the same orientation after superimposition. (D) RGA_GRAS_ is represented as a cyan surface model, and SLY1 is displayed as a purple cartoon model. The SLY1-binding surface of RGA_GRAS_ is colored yellow. (E) SHR is depicted as a blue surface model, and JKD is depicted as a teal cartoon model with the JKD-binding surface of SHR colored red. (F) Steric clash between RGA_RD_ and JKD in the superimposed structures of the RGA and SHR-JKD. The surface of JKD clashing with RGA_RD_ is colored red. (G) Cartoon model illustrating the binding interface between SLY1 and ASK2. (H, I) Surface charge distribution at the binding interface of SLY1 (H) and ASK2 (I). The electrostatic potential, ranging from red (−10 kcal/mol·*e*) to blue (+10 kcal/mol·*e*), is plotted on the solvent-accessible surfaces.

The GRAS domain of *Arabidopsis* SHORT ROOT (SHR_GRAS_) binds to the zinc-finger transcription factor JACKDAW (JKD), a member of the IDD family, through a hydrophobic surface groove^34^. When the SHR_GRAS_-JKD structure was superimposed onto that of RGA, the JKD-binding surface of SHR_GRAS_ overlapped with the SLY1-binding surface of RGA_GRAS_. Moreover, JKD exhibited steric clashes with RGA_RD_ (**Fig. 5D–5F**). This indicates that DELLA proteins bind the IDD family transcription factors in a similar manner to JKD binding to SHR_GRAS_ and that RGA_RD_ likely prevents transcription factor binding when RGA forms a complex with GA-bound GID1A.

SLY1 bound to ASK2 on the opposite surface from its RGA-binding interface through its F-box motif. This was primarily mediated by helix A3 of SLY1. Specifically, residues D48, K50, T51, A53, S56, C57, and V58 in helix A3 of SLY1 formed ionic or hydrogen bonds with ASK2 residues K137, A169, K137, N166, N166, R145 and R145, respectively (**Fig. 5G**). Additionally, SLY1-H63 in helix A4 and SLY1-R101 in helix A7 formed additional ionic bonds with ASK2-E165 (**Fig. 5G**). Like the RGA-binding surface, the ASK2-binding surface of SLY1 also showed high hydrophobicity (**Fig. 5H and 5I**). Consistent with this hydrophobicity, only ASK2 was successfully purified, but SLY1 was precipitated during purification when SLY1 and ASK2 were coexpressed (**Fig. S7J–L**).

## DISCUSSION

DELLA proteins are key regulators of plant growth and development^8–11,23^. Their activity is modulated by GA-induced GID1 binding, followed by the polyubiquitination of DELLA proteins^30–32^. In this study, we determined two cryo-EM structures of RGA complexes. The atomic models of these complexes were constructed for both the main and side chains by tracing the cryo-EM map. Notably, water molecules and GA_3_ were identified in the cryo-EM map of RGA-GID1A, despite the relatively small size of the protein complex for cryo-EM structure analysis. The atomic model in the cryo-EM structure of RGA-GID1A is approximately 86 kDa, excluding the flexible long linker. These cryo-EM structures elucidate the assembly mechanism of the ubiquitin E3 ligase responsible for the degradation of DELLA proteins. GID1A conceals GA_3_ and interacts with RGA, which in turn binds to GA_3_-bound GID1A and SLY1 through distinct binding interfaces. Additionally, SLY1 possesses separate binding interfaces for RGA and ASK2 (**Fig. 6A**). Thus, these findings reveal that the GID1A-RGA-SLY1-ASK2 complex assembles in a chain-like manner via distinct, nonoverlapping binding interfaces.

**FIGURE 6.**
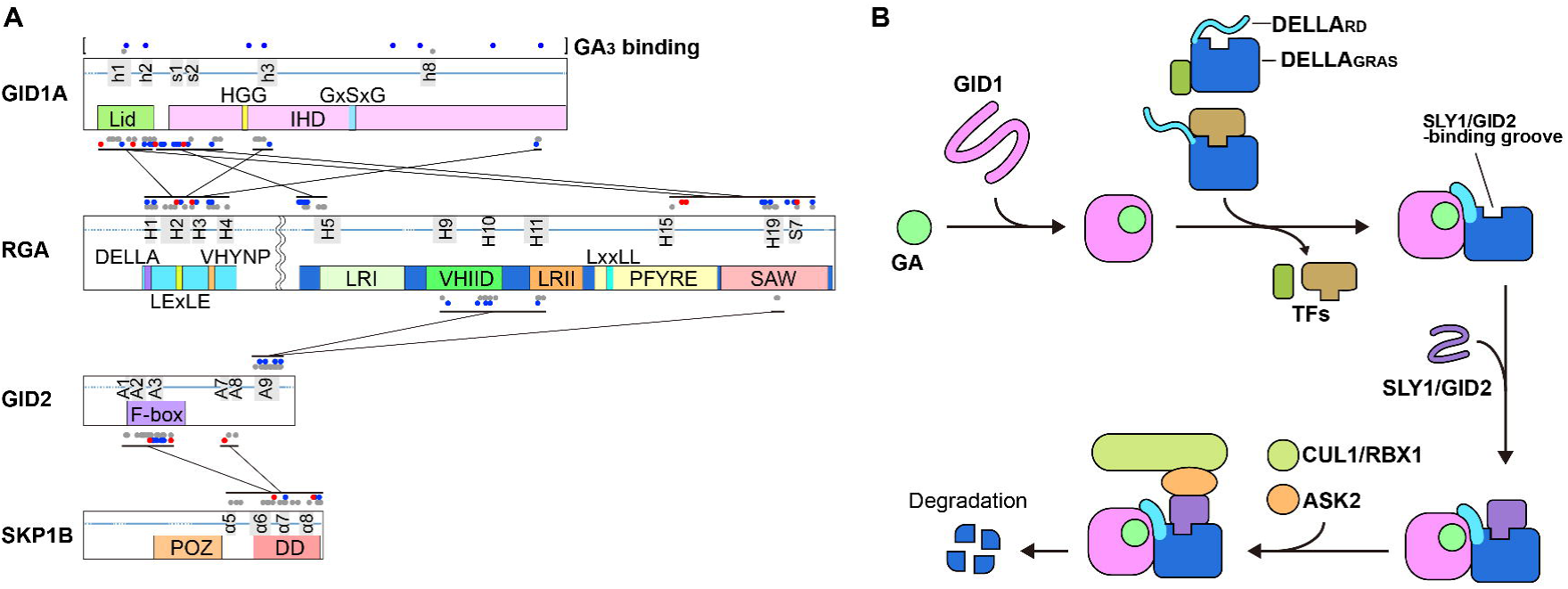
GA-induced assembly of the SCF^SLY^^1^-RGA-GID1A complex. (A) Schematic representation of the interactions between GID1A, RGA, SLY1, and ASK2. Each protein is depicted as a bar, with the lower and upper halves illustrating known structural motifs and the secondary structures at the binding interfaces. Red, blue, and gray dots indicate residues involved in ionic bonds, hydrogen bonds, and hydrophobic interactions, respectively. (B) Model of GA-induced stepwise assembly for RGA degradation and transcription factor regulation.

F-box proteins serve as substrate-binding components of SCF ubiquitin E3 ligases. The substrate binding of F-box proteins is regulated primarily by post-translational modifications of the substrate degron, such as phosphorylation and glycosylation^36^. In contrast, the substrate binding of SLY1 is regulated in response to the GA hormone^30–32^. Our cryo-EM structures elucidate the mechanism by which DELLA proteins are degraded in response to this hormone. In the cryo-EM structure of the GID1A-RGA-SLY1-ASK2 complex, SLY1 interacts with RGA_GRAS_ but not with RGA_RD_ or GID1A. Consistent with this, SLY1 is copurified with RGA_GRAS_ and ASK2 when RGA_RD_ and GID1A are absent, indicating that RGA_GRAS_ is sufficient for SLY1 binding. Therefore, the interaction between SLY1 and RGA_GRAS_ is likely regulated through RGA_RD_, as GA-bound GID1A is essential for RGA ubiquitination. The RD of DELLA proteins is stabilized upon GID1 binding through a disorder-to-order transition^6^. Consistently, RGA_GRAS_ is purified in a homogeneous form, whereas the full-length RGA tends to form aggregates. Given that RGA is costabilized by forming a heterodimer with GID1A in the presence of GA, RGA likely stabilizes through a conformational rearrangement of RGA_RD_ upon GID1 binding, thereby becoming accessible to SLY1. This suggests that DELLA proteins expose the SLY1-binding site upon GID1 binding, rendering them susceptible to ubiquitination (**Fig. 6B**). SLY1 is stabilized through its interaction with RGA. It was observed that SLY1 precipitated in the absence of RGA-GID1A, but it stabilized when copurified with RGA-GID1A and ASK2. Consistent with this observation, the C-terminal helix A9 of SLY1, which functions as a substrate-binding motif, exhibits hydrophobicity. Thus, the hydrophobic C-terminal helix of SLY1 appears to be stabilized by binding to RGA through hydrophobic interactions. Homogeneous forms of RGA and SLY1 were purified exclusively through a co-purification strategy. In summary, the results of the cryo-EM structure and protein-protein interaction analyses suggest that the SCF^SLY1^-RGA-GID1A complex is assembled in a stepwise manner, stabilizing the proteins in response to GA signals (**Fig. 6B**). In this model, GA induces the assembly of SCF^SLY1^ for the degradation of DELLA proteins. GID1_Lid_ binds GA and is costabilized with DELLA_RD_, facilitating further interactions between GID1_IHD_ and DELLA_GRAS_. The interaction between GID1 and DELLA proteins exposes the SLY1-binding surface and potentially restricts the access of transcription factors to DELLA proteins. SLY1 binds to the exposed hydrophobic groove of DELLA_GRAS_ and assembles the SCF complex for DELLA ubiquitination (**Fig. 6B**).

Some bHLH transcription factors are inactivated by binding to DELLA proteins^12–16^, whereas some IDD and bZIP family proteins are activated by binding them^17–21^. Given that the DNA-binding motifs of transcription factors are not conserved, different types of transcription factors are likely to interact with distinct surfaces of DELLA proteins. These transcription factors appear to bind to the GRAS domain of DELLA proteins^34,35^; however, the specific binding sites and regulatory mechanisms of these interactions remain unclear. Our cryo-EM structures provide insights into how DELLA proteins regulate transcription factors. A structural comparison between GID1A-RGA-SLY1 and SHR_GRAS_-JDK^34^ revealed that the zinc-finger motif of IDD family proteins accommodates the SLY1-binding groove of RGA_GRAS_ but sterically clashes with RGA_RD_. Given that the RD of DELLA proteins transitions from a random coil to an ordered conformation upon the binding of GID1A-GA ^6^, it is evident that DELLA_RD_ inhibits the binding of IDD family proteins to DELLA_GRAS_ through its interaction with GID1-GA and the resulting conformational rearrangement. In this context, the direct interaction between GID1 and DELLA_GRAS_ may also restrict the binding of other transcription factors to RGA_GRAS_, as GID1A covers a substantial surface area of RGA_GRAS_. Assuming that the binding sites of GID1 and transcription factors on DELLA_GRAS_ partially overlap, GID1 could competitively prevent transcription factors from binding to DELLA proteins (**Fig. 6B**).

## METHODS

### Plasmid preparation

The DNA sequences encoding full-length RGA (residues 1–587; UniProt ID Q9SLH3), SLY1 (residues 1–151; UniProt ID Q9STX3), and ASK2 (residues 1–171; UniProt ID Q9FHW7) were amplified from the cDNA of *A. thaliana* (Gelvin Arabidopsis CYFP-cDNA stock number CD4-58, The Arabidopsis Information Resource)^37^ via polymerase chain reaction. The gene encoding GID1A (residues 1–345; UniProt ID Q9MAA7) was synthesized following codon optimization on the basis of the codon usage preferences of *E. coli* (Bioneer, Daejeon, South Korea). The genes for *RGA* and *ASK2* were inserted into the pET-His-SUMO vector, and *SLY1* was inserted into the pRSF-His-SUMO vector^38^. Both vectors express a target protein with the N-terminal 6×His-SUMO tag removed by tobacco etch virus (TEV) protease. GID1A was inserted into the pCDF-His vector^38^, which expresses a target protein with the N-terminal 6×His removed by TEV protease.

### Purification of the GID1A-RGA complex

Plasmids for the expression of RGA and GID1A (pET-His-SUMO-RGA and pCDF-His-GID1A) were co-introduced into the *E. coli* strain BL21-Star (DE3) (Thermo Fisher Scientific, Waltham, MA, USA). The transformed cells were cultured in Luria-Bertani (LB) medium at 37 °C. When the optical density at 600 nm of the culture reached 0.6–0.7, the media was cooled at 4 °C for 1 h. Subsequently, 0.4 mM isopropyl β-thiogalactopyranoside was added to induce protein expression, along with 0.4 mM gibberellic acid (GA_3_), to promote stable expression of the RGA-GID1A complex. After being cultured at 12 °C for 36 h, the cells were harvested via centrifugation at 3,000 × *g* for 10 min, resuspended in buffer A (20 mM HEPES pH 7.5, 0.2 M NaCl, 5% (v/v) glycerol, 0.5 mM TCEP, and 0.4 mM GA_3_), and lysed via sonication. The cell lysates were treated with DNase I (Roche, Mannheim, Germany) and RNase A (Roche) (10 μg/mL each) for 30 min on ice. Insoluble debris was removed by centrifugation at 20,000 *× g* for 30 min at 4 °C.

The GID1A-RGA complex was purified using immobilized metal affinity chromatography (IMAC) and size exclusion chromatography (SEC). The clarified cell lysate was applied to a 5 mL HisTrap nickel-chelating column (Cytiva, Marlborough, MA, USA). The column was washed with 80 mM imidazole to remove nonspecifically bound proteins from the resin. Proteins bound to the resin were eluted using a linear imidazole gradient ranging from 0.08 to 1.0 M on an AKTA-purifier FPLC (Cytiva). Eluate fractions containing 6×His-SUMO-RGA and 6×His-GID1A were pooled and treated with TEV protease at 4 °C overnight. After confirming the complete cleavage of the N-terminal tags, 6×His-SUMO and 6×His, the protein solution was concentrated and loaded onto a PD-10 desalting column (Cytiva) equilibrated with buffer A to remove imidazole from the protein solution. To remove the N-terminal tag, the protein fractions eluted from the desalting column were passed through HisPur Ni-NTA resin using a gravity-flow column (Thermo Fisher Scientific, Waltham, MA, USA). The eluates were loaded onto a Superdex 200 preparative grade column (Cytiva) that had been preequilibrated with buffer B (20 mM HEPES pH 7.5, 0.2 M NaCl, and 0.5 mM TCEP). The eluate fractions corresponding to the monodisperse UV peak were pooled after confirmation via SDS-PAGE and subsequently concentrated to 5 mg/mL.

### Purification of the GID1A-RGA-SLY1-ASK2 complex

Plasmids for the expression of SLY1 and ASK2 (pRSF-His-SUMO-SLY1 and pET-His-SUMO-ASK2) were co-introduced into the *E. coli* strain BL21-star (DE3). The SLY1-ASK2 complex was expressed using the same procedures as the RGA-GID1A complex, with the exception that GA_3_ was not added to the LB medium during protein expression. The cells expressing RGA-GID1A and SLY1-ASK2 were cultured and subsequently harvested. The harvested cell pellets were combined and resuspended in buffer C (20 mM HEPES pH 7.5, 0.5 M NaCl, 5% (v/v) glycerol, 0.5 mM TCEP, and 0.4 mM GA_3_). The GID1A-RGA-SLY1-ASK2 complex was purified following the same procedures used for the RGA-GID1A complex. The final SEC was conducted using a Superdex 200 preparative grade column (Cytiva) preequilibrated with buffer D (20 mM HEPES pH 7.5, 0.5 M NaCl, and 0.5 mM TCEP). The fractions containing the GID1A-RGA-SLY1-ASK2 complex were pooled and concentrated to a final concentration of 3 mg/mL.

### Cryo-EM data collection

Purified complexes of GID1A-RGA and GID1A-RGA-SLY1-ASK2 (3 µL each) were applied to glow-discharged Cu 300 mesh QUANTIFOIL R1.2/1.3 holey carbon grids (SPI Supplies, West Chester, PA, USA). The grid was blotted using a Vitrobot Mark IV (Thermo Fisher Scientific) with humidity-saturated filter paper (Ted Pella, Redding, CA, USA). After blotting for 3 s at a blot force of 5, the grid was rapidly plunged into liquid ethane. Micrographs were collected on a 300 kV Krios G4 cryo-transmission electron microscope equipped with a Falcon 4 detector and a Selectris-X energy filter (Thermo Fisher Scientific). The energy filter slit was adjusted to 10 eV. In total, 2,362 micrographs for the RGA-GID1A complex and 1,925 micrographs for the GID1A-RGA-SLY1-ASK2 complex were collected with a pixel size of 0.7451 Å at a magnification of 165,000x, for a total dose of 60 electrons per Å^2^. The nominal defocus value ranged from −0.5 to −1.9 µm (**Table S1**).

### Data processing and structure determination

Image processing was conducted using cryoSPARC (v.4.4.0) software^39^. The raw movies were motion-corrected using Patch Motion Correction, and the defocus value for each micrograph was estimated using Patch CTF Estimation.

To construct the cryo-EM map of the GID1A-RGA complex, particles were identified using Template Picker with templates obtained from a previous pilot data collection. A total of 2,804,928 particles were picked from 2,362 micrographs and extracted with a box size of 400 pixels for 2D classification. After multiple rounds of 2D classification, a set of 273,109 particles was obtained. Particles were subsequently repicked from the micrographs using the TOPAZ program in the cryoSPARC platform^40^, yielding a total of 676,577 extracted particles. The two sets of particles were combined, and duplicates were removed. The particles were then subjected to additional rounds of 2D classification to eliminate poor-quality particles, resulting in a final selection of 507,751 particles. An initial *ab initio* model was reconstructed from these particles and classified into four classes. Of these, 265,845 particles belonging to the major class were subjected to motion correction using reference-based motion correction. Finally, the cryo-EM map of GID1A-RGA was reconstructed from the motion-corrected 251,864 particles using heterorefinement, with an estimated overall resolution of 2.66 Å (**Figs. S2 and S3**).

To reconstruct the cryo-EM map of GID1A-RGA-SLY1-ASK2, a total of 2,183,664 particles were autopicked from 1,925 micrographs using a Blob Picker, with extraction performed at a box size of 400 pixels. After multiple rounds of 2D classification, a set of 756,176 particles was obtained. The particles were then repicked from the micrographs using the TOPAZ program in the cryoSPARC platform^40^, resulting in 1,428,933 extracted particles. These two sets were combined, and duplicates were removed. The particles were then subjected to additional rounds of 2D classification to eliminate poor-quality particles and duplicates, leading to the selection of 682,192 particles. An initial *ab initio* model was reconstructed from the combined particles and classified into three classes. Of these, 277,898 particles from the major class were motion-corrected using reference-based motion correction. The cryo-EM map of GID1A-RGA-SLY1-ASK2 was reconstructed from the motion-corrected 277,150 particles using nonuniform refinement and classified into 20 clusters using 3D variability analysis. The final cryo-EM map was reconstructed using nonuniform refinement with 65,737 particles selected from a 3D variability, achieving an overall resolution of 2.80 Å. To enhance the map quality of SLY1 and ASK2, a cryo-EM map focusing on SLY1-ASK2 was reconstructed at a resolution of 3.07 Å using local refinement (**Figs. S8 and S9**). The final map was reconstructed by combining the cryo-EM map of GID1A-RGA-SLY1-ASK2 with the focused map of SLY1-ASK2 using the Phenix.combine_focused_maps^41^.

All reported resolutions were estimated using the gold-standard Fourier shell correlation (FSC) = 0.143 criterion (**Figs. S3 and S9**). The cryo-EM map processing statistics are summarized in Table S1, and the overall workflow of cryo-EM data processing is illustrated in Figs. S2 and S8.

### Atomic model building and structure analysis

The AlphaFold^42^ models of GID1A, RGA, SLY1, and ASK2 were used as templates for model building. Each model was fitted into the cryo-EM map using ChimeraX^43^, rebuilt by tracing the cryo-EM map in COOT^36^ and refined using Phenix.refinement^41,44^. Water molecules were added using Phenix.douse^41^ and subsequently confirmed manually. The statistics for Cryo-EM data collection and refinements are summarized in Table S1. Molecular interactions were analyzed using PISA^45^ and DIMPLOT^46^. Figures were generated using ChimeraX^43^, PyMOL^47^, and ALSCRIPT^48^. The atomic models and cryo-EM maps have been deposited in the Protein Data Bank with the PDB accession numbers: xxxx for GID1A-RGA and xxxx for GID1A-RGA-SLY1-ASK2.

### SEC-MALS

The molar masses of the GID1A-RGA and GID1A-RGA-SLY1-ASK2 complexes were measured using an SEC-MALS instrument (Wyatt Technology, Santa Barbara, CA, USA). Purified GID1A-RGA (100 µL, 1.5 mg/mL) and GID1A-RGA-SLY1-ASK2 (100 µL, 1 mg/mL) complexes were injected into a Superdex 200 Increase 10/300 GL column (Cytiva) equilibrated with buffer B (20 mM HEPES pH 7.5, 0.5 M NaCl, and 0.2 mM TCEP). The eluate was then applied to inline DAWN Heleos II MALS and Optilab T-Rex differential refractive index detectors (Wyatt Technology). Data analysis was performed using the ASTRA 6 software package (Wyatt Technology), and data were visualized using SigmaPlot 14.0 (Grafiti LLC, Palo Alto, CA, USA).

## ACKNOWLEDGMENTS

We would like to express our gratitude to the staff at the Korea Basic Science Institute for their assistance with SEC-MALS analyses.

## DATA AND CODE AVAILABILITY

The final coordinates and cryo-EM maps that support the findings of this study have been deposited in the Worldwide Protein Data Bank (www.wwpdb.org) with the following PDB IDs (9LUM, 9LUN, 9LUO, and 9LUP) and EMDB IDs (EMD-63398, EMD-63399, EMD-63400, and EMD-63401).

## CONFLICT OF INTEREST

The authors declare no conflict of interest.

## AUTHOR CONTRIBUTIONS

S.I. and K.P. performed the experiments. S.I., K.P., E.K., and D.Y.K. carried out the data analysis. E.K. and D.Y.K. wrote the manuscript with contributions from all other authors.

## Notes

### Competing Interest Statement

The authors have declared no competing interest.

