## Supplementary information for "Structural analyses of gibberellin-mediated DELLA protein degradation"

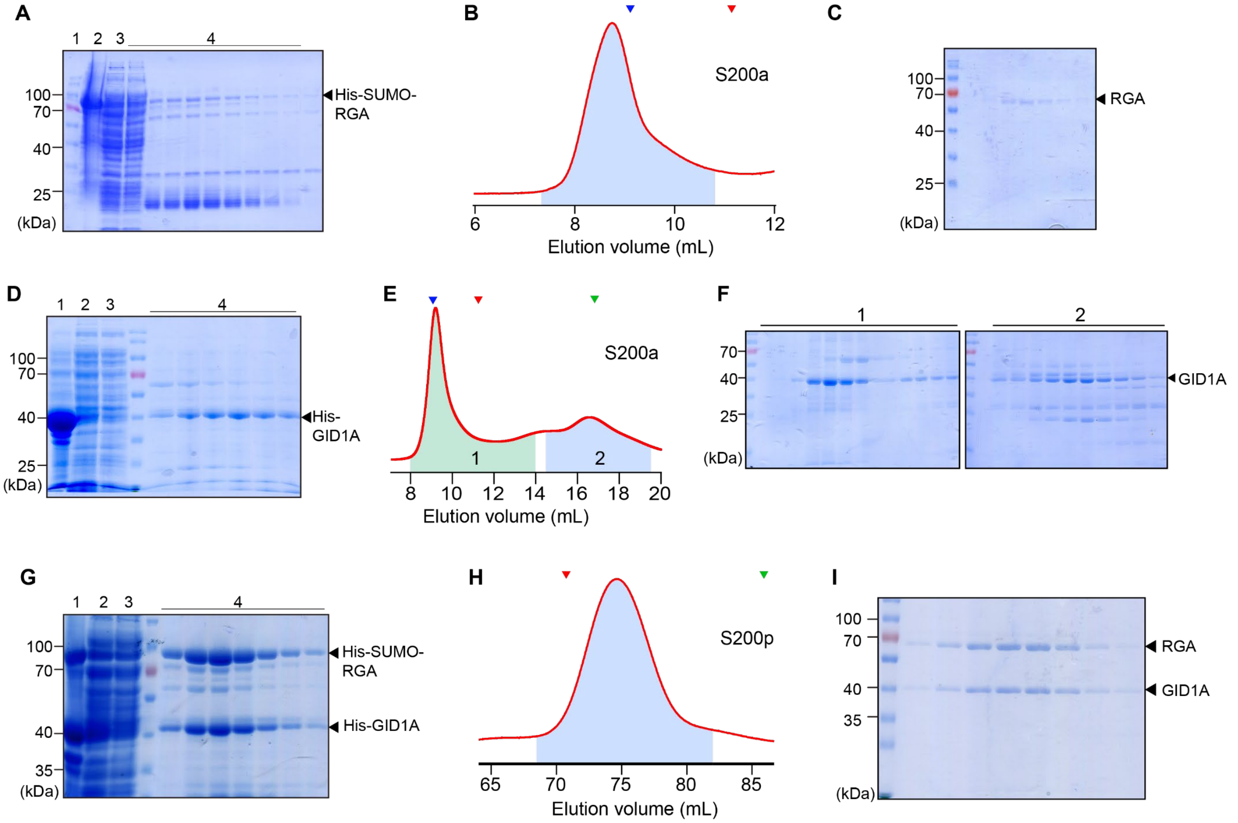


**Supplemental Figure 1. Co-purification and stability of RGA and GID1A.** RGA, GID1A, and the GID1A-RGA complex were purified in the following steps under buffer conditions containing 0.2 M NaCl: immobilized metal affinity chromatography (IMAC), N-terminal tag removal using tobacco etch virus (TEV) protease and Ni-chelating resin, followed by size-exclusion chromatography (SEC). The left, middle, and right figures indicate the SDS-PAGE of IMAC, the absorbance curve at 280 nm from SEC, and the SDS-PAGE of SEC, respectively. Lanes 1 and 2 in the left figures represent the insoluble and soluble fractions of the cell lysate after centrifugation, respectively. Lane 3 shows an unbound sample from the Ni-chelating resin, and Lane 4 shows the proteins eluted from the resin with an imidazole gradient during IMAC. The green and blue shaded area under the absorbance curve at 280 nm corresponds to the SEC fractions analyzed using SDS-PAGE. S200a and S200p indicate the Superdex 200 Increase and Superdex 200 preparative grade columns used for SEC, respectively. Blue, red, and green triangles indicate the elution volumes of ferritin (440 kDa), aldolase (158 kDa), and ovalbumin (44 kDa), which were used as molecular standards. (A–C) Purification of RGA. (D–F) Purification of GID1A expressed in the presence of $GA_3$. (G–I) Co-purification of the RGA-GID1A complex co-expressed in the presence of $GA_3$.

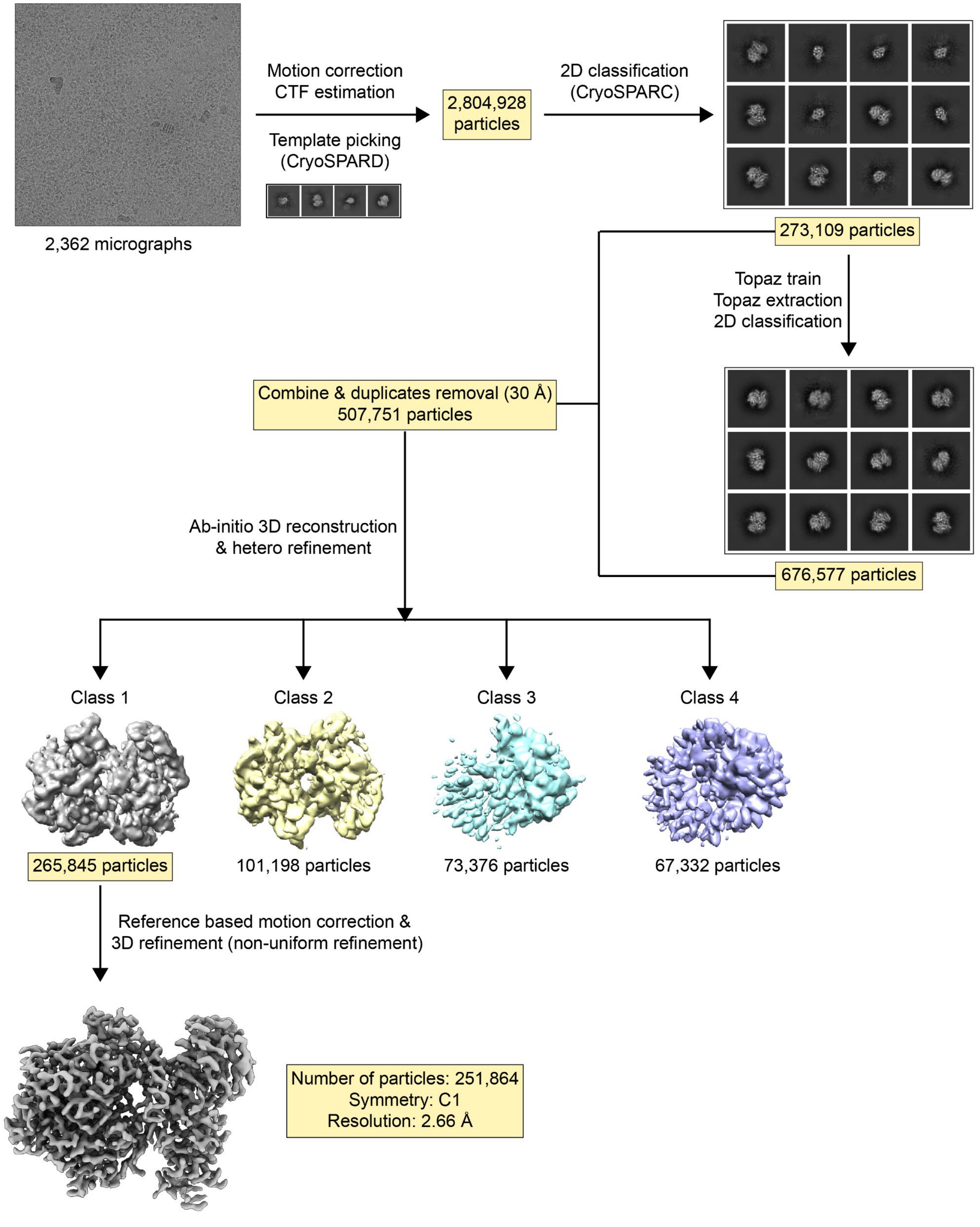


**Supplemental Figure 2. Cryo-EM data processing workflow for the GID1A-RGA complex.**

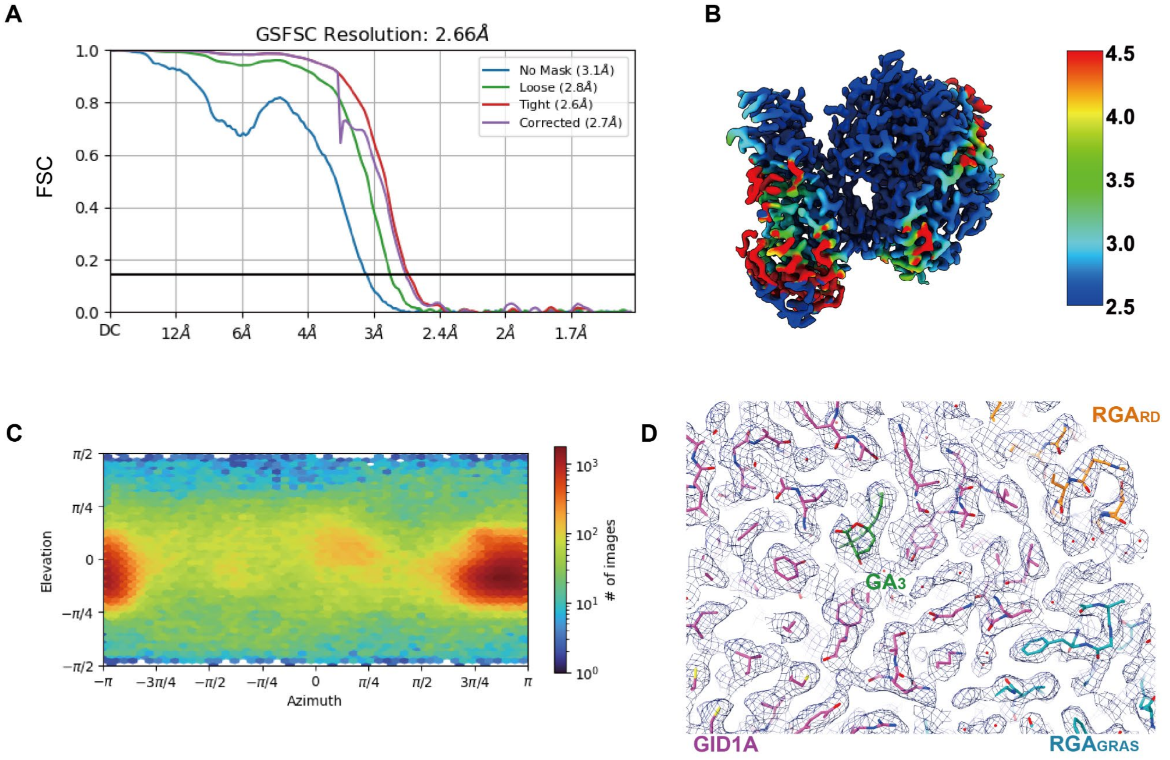


**Supplemental Figure 3. Cryo-EM data analysis of GID1A-RGA.** (A) Fourier shell correlation (FSC) curves, where the overall resolution was estimated at an FSC value of 0.143. (B) Cryo-EM map of RGA-GID1A showing the local resolution. (C) Angular distribution of particle projections for GID1A-RGA. (D) Cryo-EM map of RGA-GID1A superimposed with its atomic model.

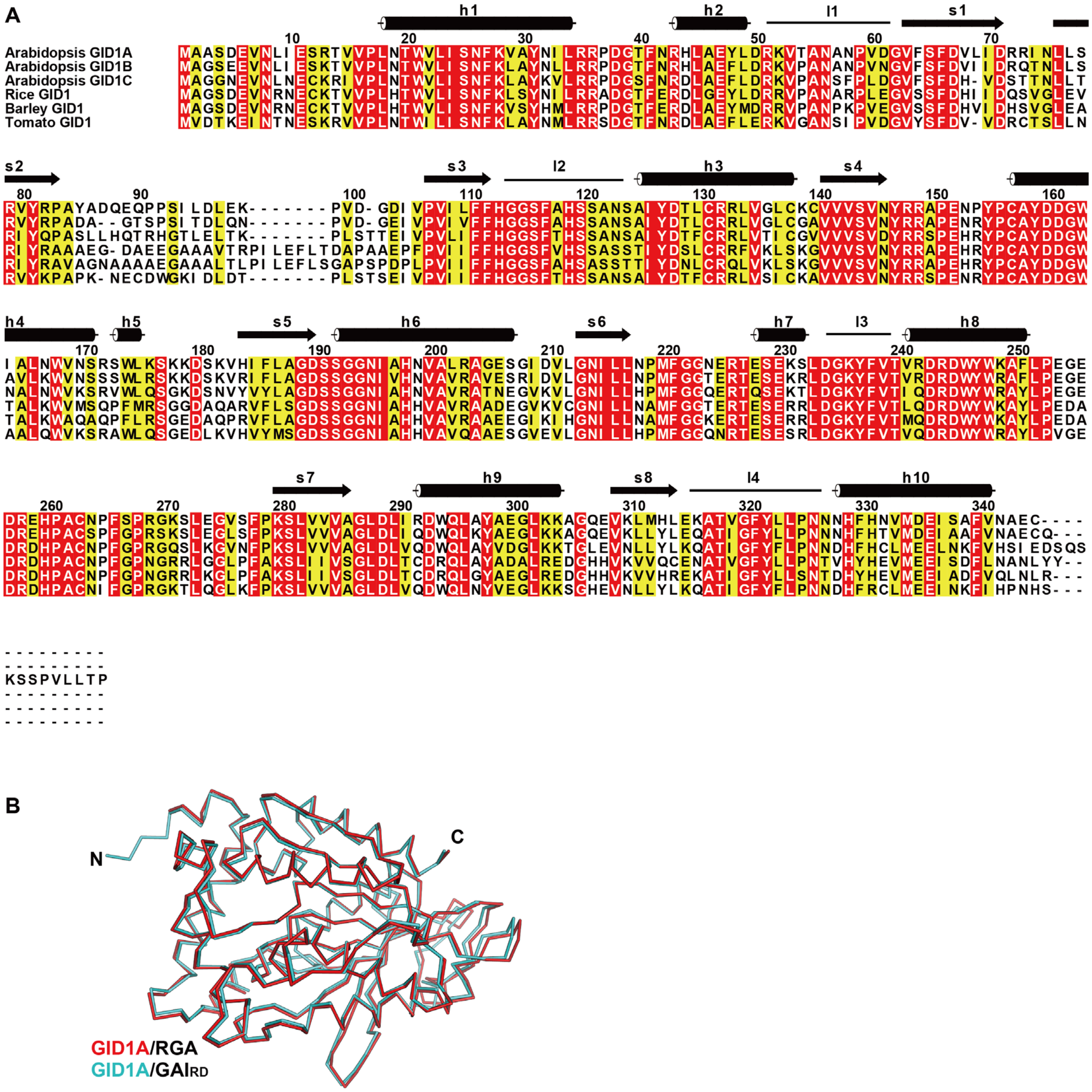


**Supplemental Figure 4. Structural analysis of GID1A.** (A) Sequence alignment of GID1A and its homologs. Black cylinders and arrows represent the helices and strands of GID1A, respectively. Blue lines indicate loops described in the main text. (B) Structural comparison of GID1A complexed with RGA and RAIRD.

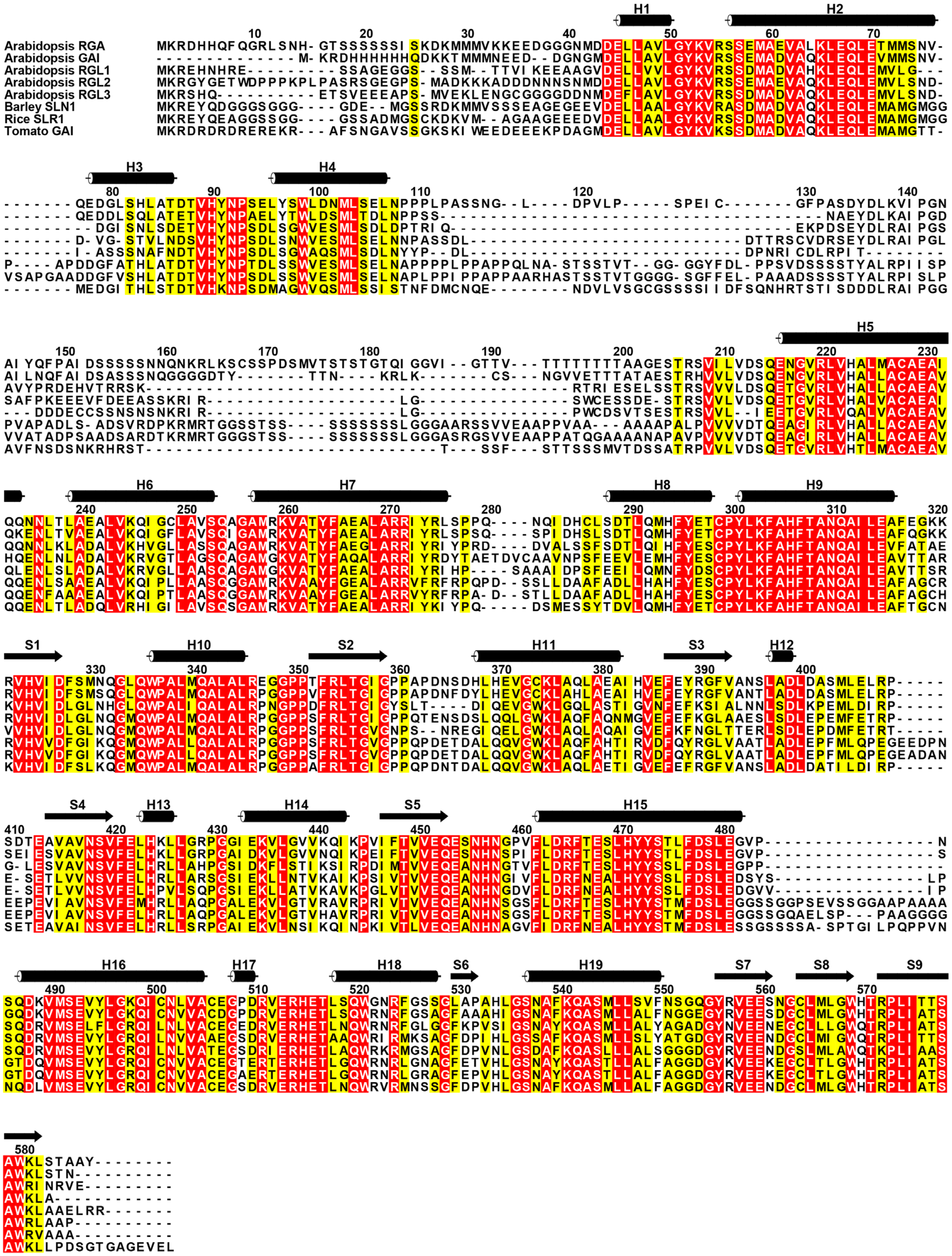


**Supplemental Figure 5. Sequence alignment of DELLA proteins.** Black cylinders and arrows represent the helices and strands of RGA, respectively.

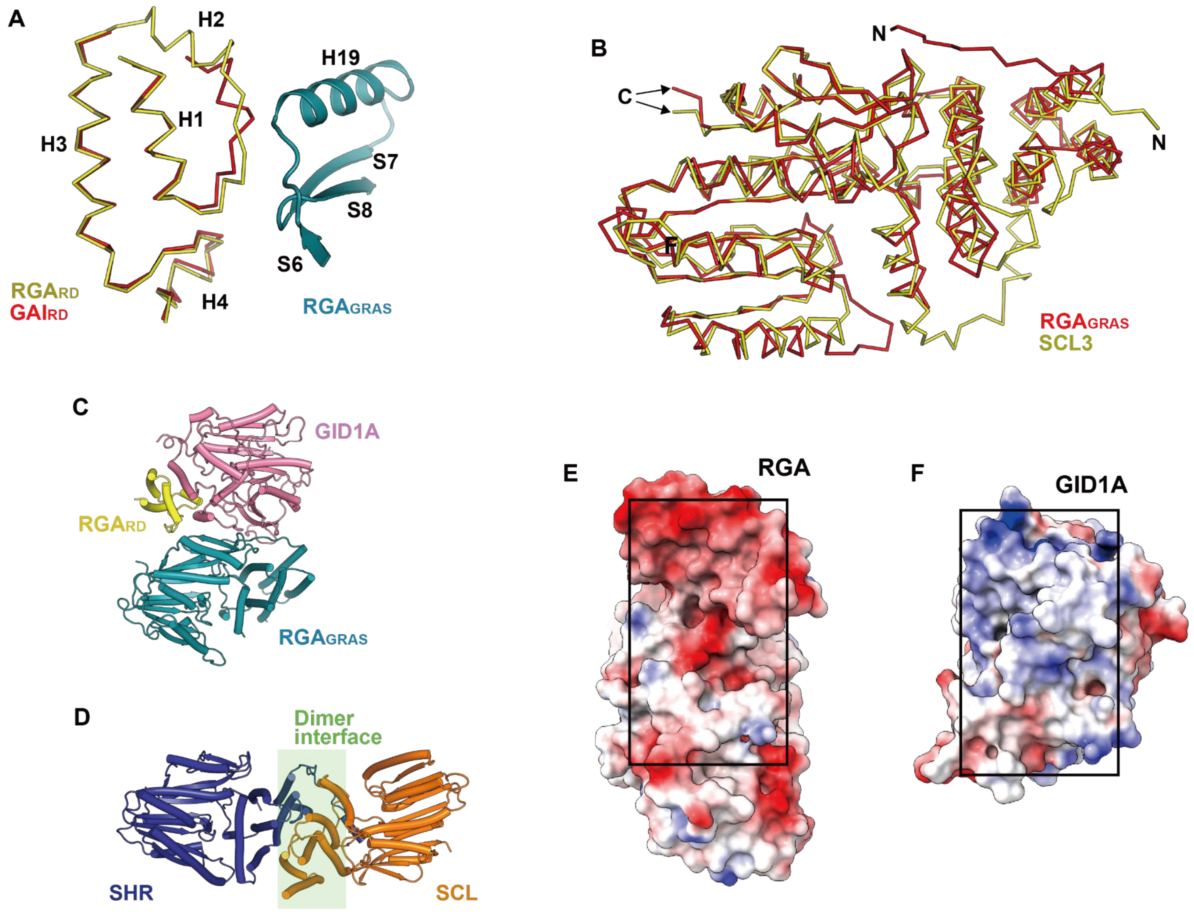


**Supplemental Figure 6. Structure and interaction analysis of RGA.** (A) Structural comparison of $RGA_{RD}$ and $GAI_{RD}$. $RGA_{RD}$ from the cryo-EM structure of GID1A-RGA and $GAI_{RD}$ from the crystal structure of GID1A-$GAI_{RD}$ (PDB ID: 2ZSH) are superimposed and depicted as Cα trace models in yellow and red, respectively. $RGA_{GRAS}$ at the interaction interface with $RGA_{RD}$ is shown as a cyan cartoon model. (B) Structural comparison of $RGA_{GRAS}$ and SCL3 (PDB ID: 6KPD). Superimposed $RGA_{GRAS}$ and SCL3 are shown as Cα trace models in red and yellow, respectively. (C, D) Dimer interface of GRAS domains. $RGA_{GRAS}$ from the cryo-EM structure of RGA-GID1A is superimposed onto SHR from the crystal structure of SHR-SCL. RGA-GID1A (C) and SHR-SCL (D) are shown in the same orientation after superimposition. The dimer interface of SHR-SCL is highlighted with a green background. (E, F) Surface charge distribution on the binding interfaces of RGA (E) and GID1A (F). Electrostatic potential, ranging from red (−10 kcal/mol·*e*) to blue (+10 kcal/mol·*e*), is plotted on the solvent-accessible surfaces.

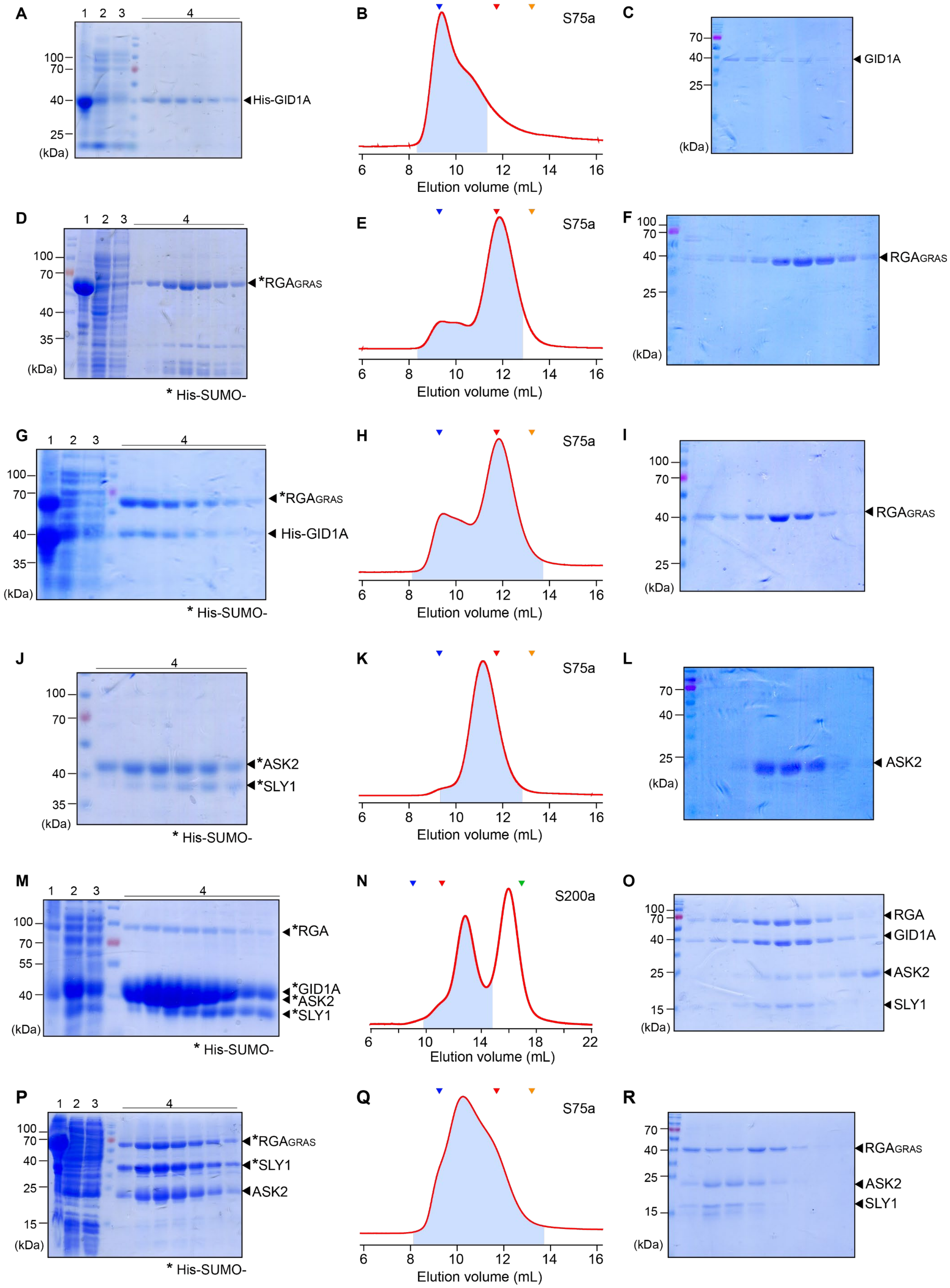
A
1 2 3 4
100
70
40
25
(kDa)
His-GID1A
B
S75a
6 8 10 12 14 16
Elution volume (mL)
C
70
40
25
(kDa)
GID1A
D
1 2 3 4
100
70
40
35
(kDa)
*RGAGRAS
* His-SUMO-
E
S75a
6 8 10 12 14 16
Elution volume (mL)
F
100
70
40
25
(kDa)
RGAGRAS
G
1 2 3 4
100
70
40
35
(kDa)
*RGAGRAS
His-GID1A
* His-SUMO-
H
S75a
6 8 10 12 14 16
Elution volume (mL)
I
100
70
40
25
(kDa)
RGAGRAS
J
4
100
70
40
35
(kDa)
*ASK2
*SLY1
* His-SUMO-
K
S75a
6 8 10 12 14 16
Elution volume (mL)
L
70
40
25
(kDa)
ASK2
M
1 2 3 4
100
70
55
40
(kDa)
*RGA
*GID1A
*ASK2
*SLY1
* His-SUMO-
N
S200a
6 10 14 18 22
Elution volume (mL)
O
100
70
40
25
15
(kDa)
RGA
GID1A
ASK2
SLY1
P
1 2 3 4
100
70
40
25
15
(kDa)
*RGAGRAS
*SLY1
ASK2
* His-SUMO-
Q
S75a
6 8 10 12 14 16
Elution volume (mL)
R
70
40
25
15
(kDa)
RGAGRAS
ASK2
SLY1

**Supplemental Figure 7. Co-purification and stability of the GID1A-RGA-SLY1-ASK2 complex.** All proteins were purified under high-salt buffer conditions containing 0.5 M NaCl, following these steps: IMAC, removal of the N-terminal tag using TEV protease and Ni-chelating resin, and SEC. The left, middle, and right figures indicate the SDS-PAGE of IMAC, the absorbance curve at 280 nm from SEC, and the SDS-PAGE of SEC, respectively. Lanes 1 and 2 in the left figures represent the insoluble and soluble fractions of the cell lysate obtained after centrifugation, respectively. Lane 3 shows the sample unbound to the Ni-chelating resin, and Lane 4 shows the proteins eluted from the resin with an imidazole gradient during IMAC. The blue shaded area under the absorbance curve at 280 nm was analyzed using SDS-PAGE during SEC. S75a and S200a indicate the Superdex 75 Increase and Superdex 200 Increase columns used for SEC, respectively. Blue, red, green, and orange triangles indicate the elution volumes of ferritin (440 kDa), aldolase (158 kDa), ovalbumin (44 kDa), and carbonic anhydrase (29 kDa), which were used as molecular standards. (A–C) Purification of GID1A. (D–F) Purification of $RGA_{GRAS}$. (G–I) Purification of $RGA_{GRAS}$ and GID1A co-expressed in the presence of $GA_3$. GID1A precipitated during purification. (J–L) Purification of co-expressed SLY1 and ASK2. SLY1 precipitated during purification. (M–O) Co-purification of GID1A-RGA-SLY1-ASK2. GID1A-RGA and SLY1-ASK2 were separately expressed and co-purified by mixing cell lysates. The four proteins co-eluted during SEC. (P–R) Co-purification of $RGA_{GRAS}$, SLY1, and ASK2. $RGA_{GRAS}$ and SLY1-ASK2 were separately expressed and co-purified by mixing cell lysates. SLY1 was stabilized during co-purification with $RGA_{GRAS}$ and ASK2.

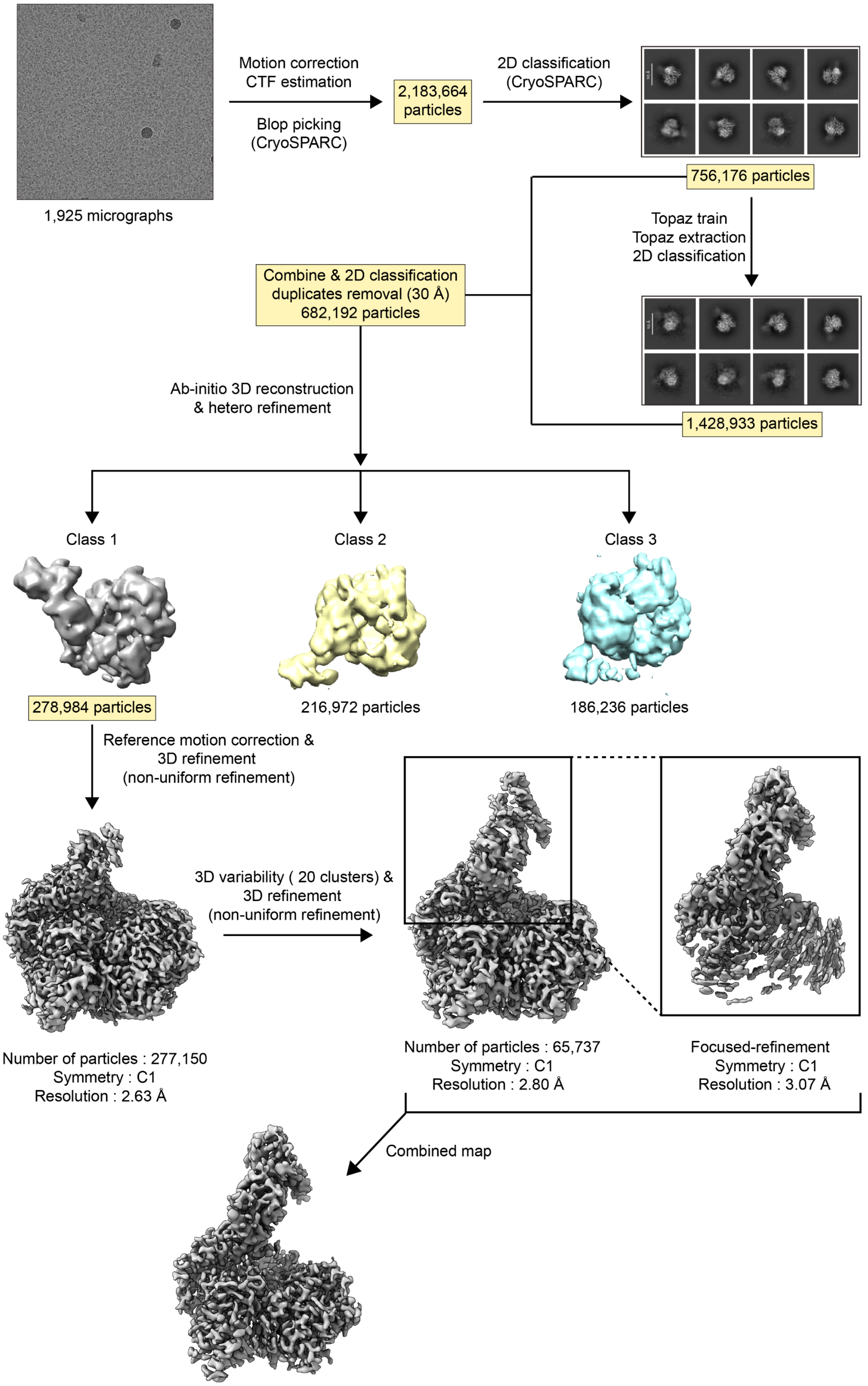


**Supplemental Figure 8. Cryo-EM data processing workflow for the GID1A-RGA-SLY1-ASK2 complex.**

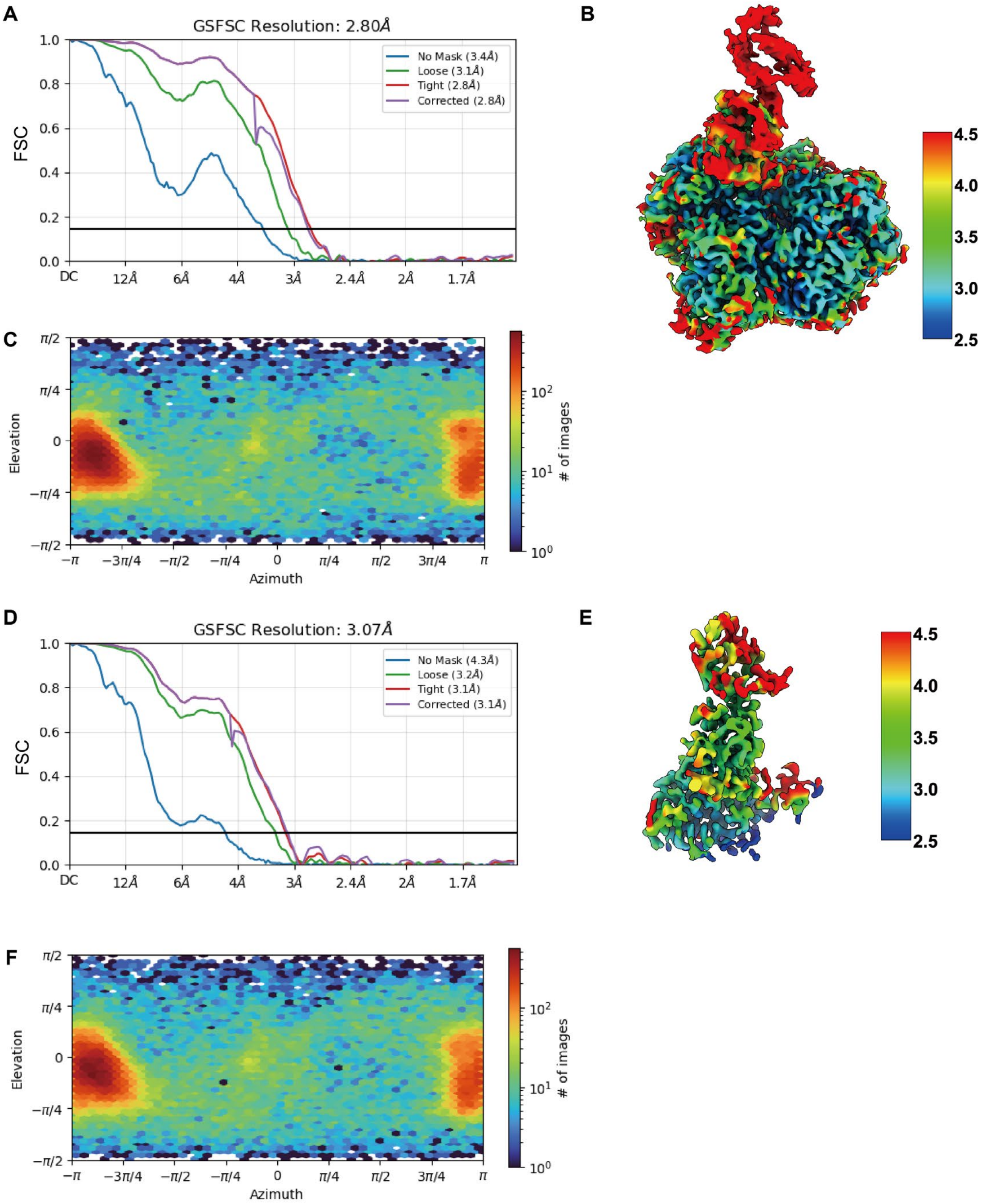


**Supplemental Figure 9. Cryo-EM data analysis of the GID1A-RGA-SLY1-ASK2 complex.** (A–C) Analysis of the GID1A-RGA-SLY1-ASK2 Cryo-EM map. (A) FSC curves. The overall resolution was estimated based on the FSC threshold of 0.143 (horizontal line). (B) Cryo-EM map displaying the local resolution, ranging from 2.5 Å to 4.5 Å, shown using a rainbow color spectrum on the cryo-EM map. (C) Angular distribution of particle projections for GID1A-RGA-SLY1-ASK2. (D–F) Analysis of the cryo-EM map, focusing on SLY1-ASK2. (D) FSC curves. (E) Cryo-EM map displaying the local resolution. (F) Angular distribution of particle projections.

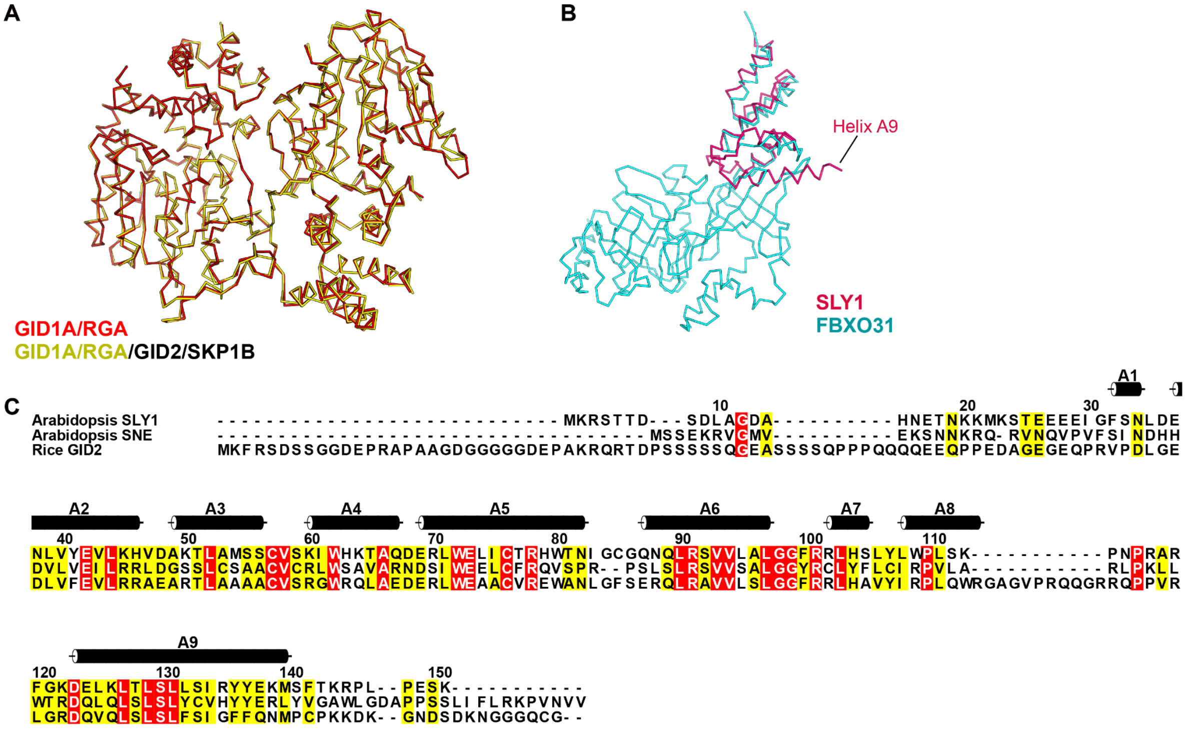


**Supplemental Figure 10. Structural analysis of the GID1A-RGA-SLY1-ASK2 complex.** (A) Structural comparison of GID1A-RGA from the cryo-EM structures of GID1A-RGA and GID1A-RGA-SLY1-ASK2. (B) Structural comparison between SLY1 and FBXO31. SLY1 and FBXO31 (PDB ID: 5VZT) are superimposed and depicted as Cα trace models in red and cyan, respectively. (C) Sequence alignment of SLY1 and its homologs. Black cylinders and arrows represent the helices and strands of SLY1.

**Supplemental Table 1. Data-collection and refinement statistics for the structure determination**

| **Data set** | GID1A-RGA | GID1A-RGA-SLY1-ASK2 |
|---|---|---|
| **Data collection** | | |
| Microscope | Titan Krios G4 | Titan Krios G4 |
| Detector | Falcon4 | Falcon4 |
| Magnification | 165,000 | 165,000 |
| Voltage (kV) | 300 | 300 |
| Electron Exposure ($e^-/Å^2$) | 60 | 60 |
| Defocus Range (μm) | -0.5 to -1.9 | -0.5 to -1.9 |
| Pixel size (Å) | 0.7451 | 0.7451 |
| No. frames/movie | 60 | 60 |
| Total exposure time (sec) | 5.32 | 5.24 |
| Number of movies | 2,362 | 1,925 |
| Symmetry imposed | *C*1 | *C1* |
| No. initial particles | 2,804,928 | 2,183,664 |
| No. final particles | 251,864 | 65,737 |
| Map resolution (Å) | 2.66 | 2.80 |
| FSC threshold | 0.143 | 0.143 |
| **Model refinement** | | |
| Composition | | |
| Atoms | 6,321 | 8.088 |
| Residues | 772 | 1,021 |
| Water | 246 | |
| Ligands | $GA_3$: 1 | $GA_3$: 1 |
| Bonds (RMSD) | | |
| Length (Å) | 0.010 | 0.007 |
| Angle (°) | 1.147 | 0.694 |
| Mean B-factors | | |
| Protein | 26.78 | 51.77 |
| Water | 25.54 | |
| Ligand | 17.77 | 35.22 |
| Ramachandran plot (%) | | |
| Favored | 96.60 | 95.82 |
| Allowed | 3.40 | 4.18 |
| Outliers | 0.00 | 0.00 |
| Rotamer outliers (%) | 1.83 | 2.84 |
| Cβ outliers (%) | 0.00 | 0.00 |
| Clash score | 4.31 | 8.08 |
| MolProbity score | 1.63 | 2.07 |
| **Model vs. Data** | | |
| CC (mask) | 0.84 | 0.77 |
| CC (box) | 0.71 | 0.67 |
| CC (peaks) | 0.70 | 0.63 |
| CC (volume) | 0.79 | 0.73 |
| Mean CC for ligands | 0.66 | 0.67 |
